# Transcriptomic profiling of healthy individual-derived LCLs revealed inter-individual variability towards-hypoxia-responsive pathways

**DOI:** 10.1101/2024.08.06.606933

**Authors:** Dayanidhi Singh, Komal Mehta, Ritu Rani, Satyam Kumar Agrawal, Bhavana Prasher

**Affiliations:** Centre of Excellence for Applied Development of Ayurveda Prakriti and Genomics, CSIR-Institute of Genomics & Integrative Biology, Delhi, India; CSIR’s Ayurgenomics unit TRISUTRA (Translational Research & Innovative Science ThRough Ayurgenomics), CSIR-Institute of Genomics & Integrative Biology, Delhi, India; Genomics & Molecular Medicine, CSIR-Institute of Genomics & Integrative Biology, Delhi, India; Academy of Scientific and Innovative Research, Ghaziabad, Uttar Pradesh, India

**Keywords:** Hypoxia, Lymphoblastoid Cell Lines, Transcriptomics, Prakriti, Inter-individual Variability

## Abstract

Hypoxia, or low oxygen levels, affects various developmental, physiological, and pathological processes. It’s been consistently reported that there is an inter-individual variability at genetic and molecular pathways related to oxygen sensing and response. Understanding the underlying variability towards hypoxia sensing and response in health and disease conditions is challenging. The *Prakriti* stratification method of Ayurveda offers solutions, which classifies healthy individuals into different groups based on multisystem phenotyping. Our lab has previously used this method and provided evidence for the variability in hypoxia responsiveness physiologically among healthy individuals at population levels.

Our current study seeks to understand hypoxia sensing and response pathways at cellular levels. We used eight Lymphoblastoid cell lines (LCLs) developed from healthy individuals of extreme *Prakriti* types. Hypoxia challenge experiments were performed using 0.2% oxygen for 24 and 48 hrs. of chronic hypoxia and captured global transcriptomics profiles. Differentially expressed genes revealed activation of core hypoxia-induced transcriptomic signatures, such as HIF-1⍺ signaling and their metabolic reprogramming in pooled as well as in all Prakriti groups. However, there were *Prakriti*-specific differences, such as activation of TGF-β mediated ROS and PI3K/AKT/mTOR driven mTORC^1^ complex in Kapha, downregulation of cholesterol homeostasis and regulation of Phosphoinositide biosynthesis in membrane potential observed in the Pitta group. In contrast, ER stress-induced activation of cell survival response via Unfolded protein response in the Vata group. The *Prakriti* stratification method will provide a novel method to understand inter-individual differences in hypoxia response pathways.

**Significance:** Hypoxia can significantly impact various aspects of our health and well-being. All nucleated cells sense and respond to hypoxia, depending upon their cellular and metabolic activities. Its wide utility and spatiotemporal regulation make it a crucial target to study. We have used the *Prakriti* stratification method of Ayurveda to explore hypoxia sensing and response at cellular levels. Lymphoblastoid Cell Lines (LCLs) developed from the peripheral blood of stratified healthy individuals have been utilized to study the expression level variability at the baseline and hypoxia-induced conditions. The outcomes of our study will be crucial to understanding inter-individual variability in response to hypoxia overlayed baseline variations. Resulting in differential susceptibility towards hypoxic response in health and contributes to understanding variable outcomes in disease conditions. Molecular targets from our study will further be utilized for interventional drug targets in hypoxia-induced disease conditions.

## Introduction

Oxygen is essential to our well-being as it is a critical component for cellular respiration, which produces the energy needed for all cellular and physiological processes in our body. The proper functioning of our cellular system requires optimum oxygen concentrations. Eukaryotes have developed mechanisms to sense and respond to oxygen level variations throughout evolution(1). Even within an organism, each cell type has its own oxygen requirements depending upon its cellular and metabolic activity. Low oxygen levels, also known as hypoxia, can affect a wide range of biological processes with different outcomes in different contexts, such as developmental, physiological, and pathological conditions(2). Significant developments in hypoxia biology have occurred in the last few decades. These developments have led to a better understanding of the essential genes involved in sensing and downstream signaling to hypoxia in healthy and diseased states(3). One of the most important genes is the Hypoxia-Inducible Factor alpha gene (*HIF-α*), which is expressed in hypoxic conditions and its activity is regulated by the oxygen-dependent Prolyl hydroxylase domain 2 (*PHD2*) also known as Egg Laying Nine-1 (*EGLN1*) (4). In hypoxic conditions, *PHD2* becomes inactive, leading to the stabilization of the HIF-α protein in the cytoplasm and translocation to the nucleus, where it forms heterodimer with its ubiquitously expressed HIF-ꞵ subunit and binds to the HRE (Hypoxia Response Element) transcription sites to activate hypoxic response. Activated HIF (Hypoxia-Inducible Factor) can further initiate the transcriptional cascade of genes to generate a hypoxic response that promotes cell survival. It can initiate metabolic reprogramming by promoting glycolysis over the oxygen-consuming TCA cycle(5). It can also activate vascular remodelling and angiogenesis genes, to enhance the supply of oxygen through blood(6). HIF can directly bind and activate histone modifiers, leading to the activation and suppression of target genes(7). For instance, hypoxia-mediated suppression of cell proliferation, protein synthesis, and regulation of immune processes are widely reported in multiple studies. Under chronic hypoxic conditions, cells try to promote survival by activating autophagy(8). However, if hypoxia persists, cells will ultimately undergo apoptosis (9) (Fig:1).

**Fig 1.**
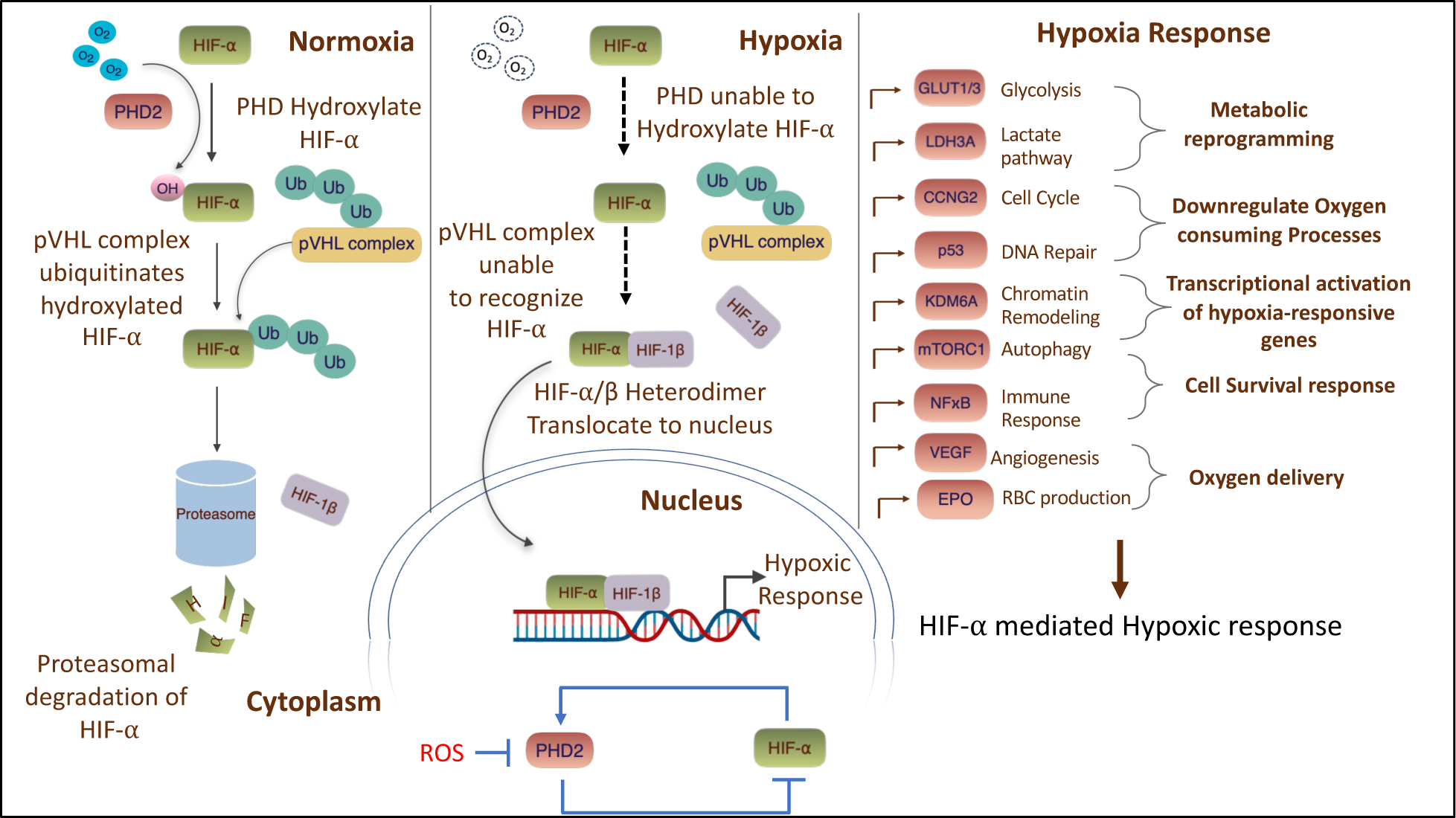
Overview of hypoxia sensing and response pathways

The above-mentioned cellular signaling pathways impact the regulation of oxygen levels in the body, and any deviation to these pathways can affect the outcomes in physiological and pathological conditions. For instance, enhanced glycolysis in high altitude conditions favours adaptation through altered metabolism, but the same can also allow cancer cells to survive and proliferate in low oxygen conditions and contribute to the severity of the disease. Further, variability in hypoxia levels also contributes to variability in disease progression and severity as well as response to therapy. Elevated levels of tumor hypoxia can cause resistance to radiotherapy, but this can be addressed by incorporating hypoxia-modifying therapy to enhance clinical results(10). Nevertheless, not all patients respond equally to the same hypoxia-modifying therapy. Moreover, the variability in spatial and temporal tumor hypoxia has been observed to contribute to the diversity in treatment responses to cancer therapy(10).

Genetic variability in oxygen sensing and its downstream pathways genes contributes to inter-individual variability in response to environmental and cellular hypoxic conditions. It would be important to understand the impact of baseline variability on cellular response pathways of hypoxia and its downstream consequences. Our lab has previously provided evidence of variability in hypoxia responsiveness among healthy individuals classified into basic constitution types using the Ayurvedic methods of *Prakriti* stratification(11–13).

In this study, young healthy individuals residing at sea level, when classified into *Prakriti* groups, showed genetic and gene expression variations in the *EGLN1* gene. Individuals of *Pitta* and *Kapha Prakriti* differed significantly <u>at</u> the genotypic level for rs479200 and rs480902. TT genotype of rs479200 was found to be over-represented in *Kapha* and was linked to higher *EGLN1* expression. The same *Pitta* and *Kapha*-associated variant were present in natives of high-altitude (C allele of rs479200) and High-altitude pulmonary edema (HAPE) patients (T allele of rs479200), respectively. Observations from the Indian Genome Variation Consortium(IGVC) and Human Genome Diversity Panel-Centre d’Etude de Polymorphisme Humain (HGDP-CEPH) have revealed disparate genetic lineages at high altitudes share the same ancestral allele (T) of rs480902 of *EGLN1* gene, which is overrepresented in Pitta types and positively correlated with high altitude adaptation(12).

In order to study (a) the cellular level differences amongst healthy individuals of extreme *Prakriti* and (b) the effect of environmental stress on them, we developed Lymphoblastoid Cell Lines (LCLs) from PBMCs of extreme *Prakriti* (*Vata Pitta* and *Kapha*) individuals. Using *Prakriti*-derived cellular models, we have shown that extreme *Prakriti* individuals have baseline variability in cell proliferation rates, such that *Kapha* cell lines have slower proliferation rates than *Vata* and *Pitta* cell types. However, when these cell lines are exposed to UV stress, *Vata* cell lines, one of the fast-growing LCLs, also showed higher cell death but recovered their cell number due to high inherent cell proliferation rates. Similarly, *Kapha* cell lines have slow cell proliferation rates and have less cell death after UV stress(14). These differences are vital to understanding cellular-level baseline variability and their possible contributions to response in healthy and disease conditions.

In our current study, our objective was to study the transcriptomic profile of extreme *Prakriti*-specific LCLs at baseline and hypoxic conditions at different time points to identify the molecular markers linked to the differential adaptability and outcomes of hypoxia. We have used the same eight LCLs that we developed in the earlier study. Hypoxia challenge experiments were performed using 0.2% oxygen for 24 and 48 hrs. of hypoxia, and global transcriptomics profiles were captured. We further validated microarray observations using qRT-PCR of identified differentially expressed genes. Functional validation of *Prakriti*-specific observations was performed using cellular assays. The observed differences could have a bearing on differential outcomes if observed in disease conditions and lead to the development of predictive signatures.

## Results

### Hypoxia Generation and Time Point Validation

We used hypoxia-specific markers HIF-1⍺ and EGLN1 at transcriptional and translational levels to confirm hypoxia generation at 0, 2, 4, 6, 12, 24, 48, and 72 hrs. time points using 0.2% oxygen concentration over LCLs. We have observed HIF-1⍺ activation after 4 hrs. of hypoxia, although significant activation was observed after 72 hrs. (Fig S2). However, we observed more than 60% cell death at 72 hrs. time point. In contrast, *EGLN1* levels were consistently downregulated from 2 hrs. onwards in hypoxic conditions (Fig: S2). At the protein level, EGLN1 protein was activated after 12 hrs. onwards in hypoxic conditions (Fig: S1). However, HIF-1α protein levels were inconsistent at selected time points. We noted prominent non-specific bands even with monoclonal antibodies, using 3 to 4 different types of HIF antibodies, from hydroxy HIF to HIF-1α, but could not capture the desired protein bands. Nevertheless, increased expression of EGLN1 at protein levels from 12 to 72 hrs. in hypoxic conditions suggests the activation of hypoxia-specific downstream signaling initiating after 12 hrs. (Fig: S1). After observing transcriptional and translational levels of hypoxia-specific markers, we noted the activation of the *EGLN1* gene after 24 and 48 hrs. at both levels in all LCLs (Fig: S1 and S2). Therefore, to study the global transcriptomic profiling of the hypoxic response, we selected 24 and 48 hrs. of hypoxia challenge experiments for microarray analysis.

### Hypoxia microarray experiment

We challenged all eight LCLs (3 Kapha, 2 Pitta, and 3 Vata) at 0.2% oxygen concentration for 24 and 48 hrs to explore the global transcriptomic response to hypoxia. Raw data has been submitted to GEO (GSE235776). Hierarchical clustering using intensity values of significant genes (conditional FDR F-test <0.001) were performed to visualize the overall expression pattern and identified two main clusters: one containing all normoxia and 0 hrs. control samples and the other containing all hypoxia-treated samples (with a few exceptions) few normoxia samples from *Vata* and *Kapha* at 24 hrs. and *Kapha* at 48 hrs. showed different clustering patterns (Fig: 2).

**Fig 2:**
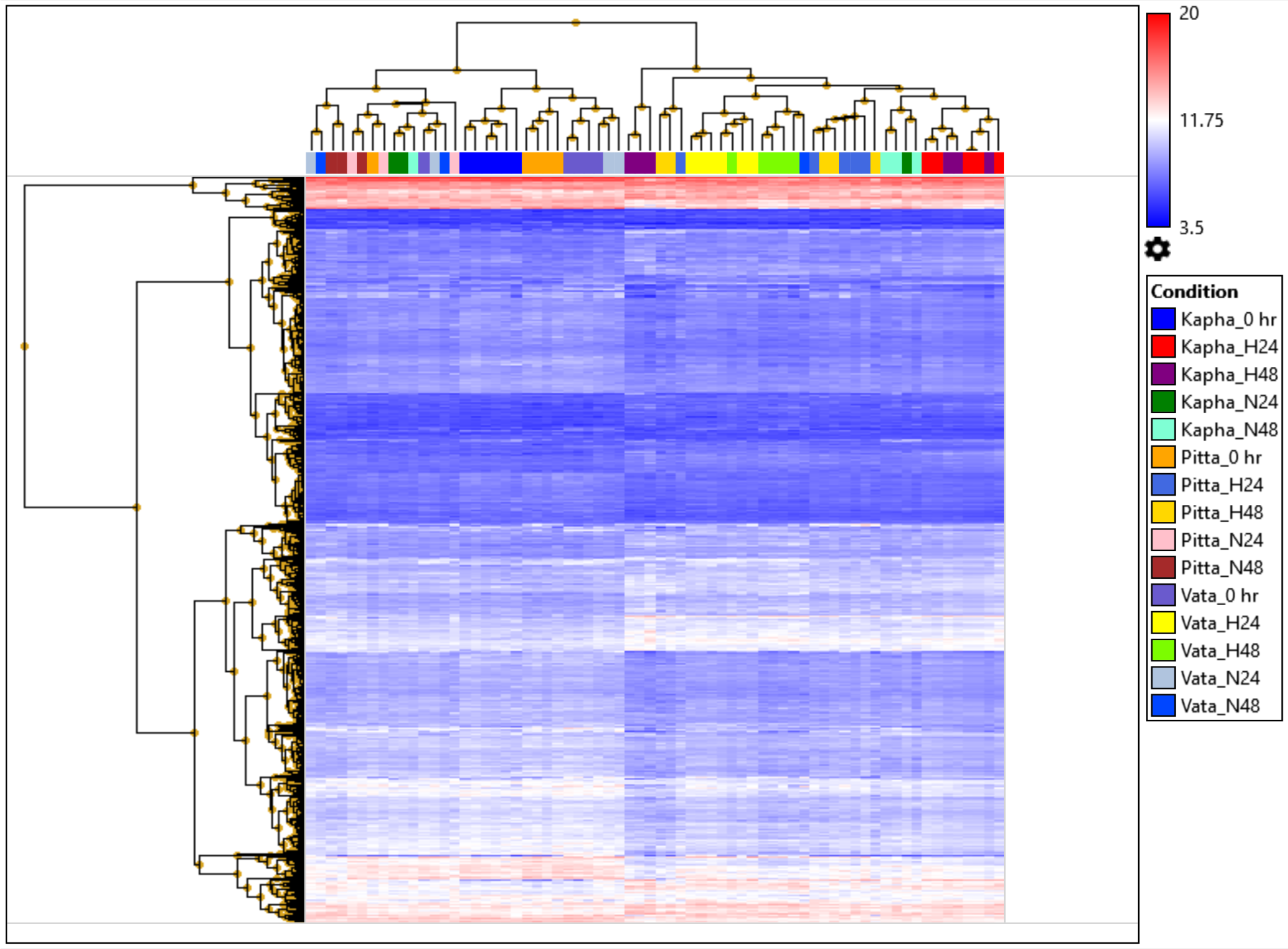
Hierarchical clustering of all 68 microarray samples (Condition FDR F-Test <0.05). Nomenclature 0 hrs. (untreated control), H24 (Hypoxia 24 hrs.), H48 (Hypoxia 48 hrs.), N24 (Normoxic 24 hrs.) and N48 (Normoxic 48 hrs.)

### Baseline Transcriptomic differences in extreme *Prakriti* LCLs

PCA plot and hierarchical clustering of baseline transcriptomic data of all three extreme *Prakriti* cell lines show segregation in separate clusters (Fig: 3). We compared the 0 hr. control samples of Vata-Pitta and Kapha among themselves to identify baseline expression differences. In the Kapha vs Pitta comparison, 2850 genes, Kapha vs Vata 2661, and Pitta vs Vata 2630 genes were differentially expressed at p-value (<0.05) significance. Out of these DEGs, 107 genes were common among all three comparisons. Differentially Expressed Genes (DEGs) were analyzed for pathway enrichment using GSEA (Gene Set Enrichment Analysis) and IPA analysis using p-value <0.05 significance.

**Fig 3:**
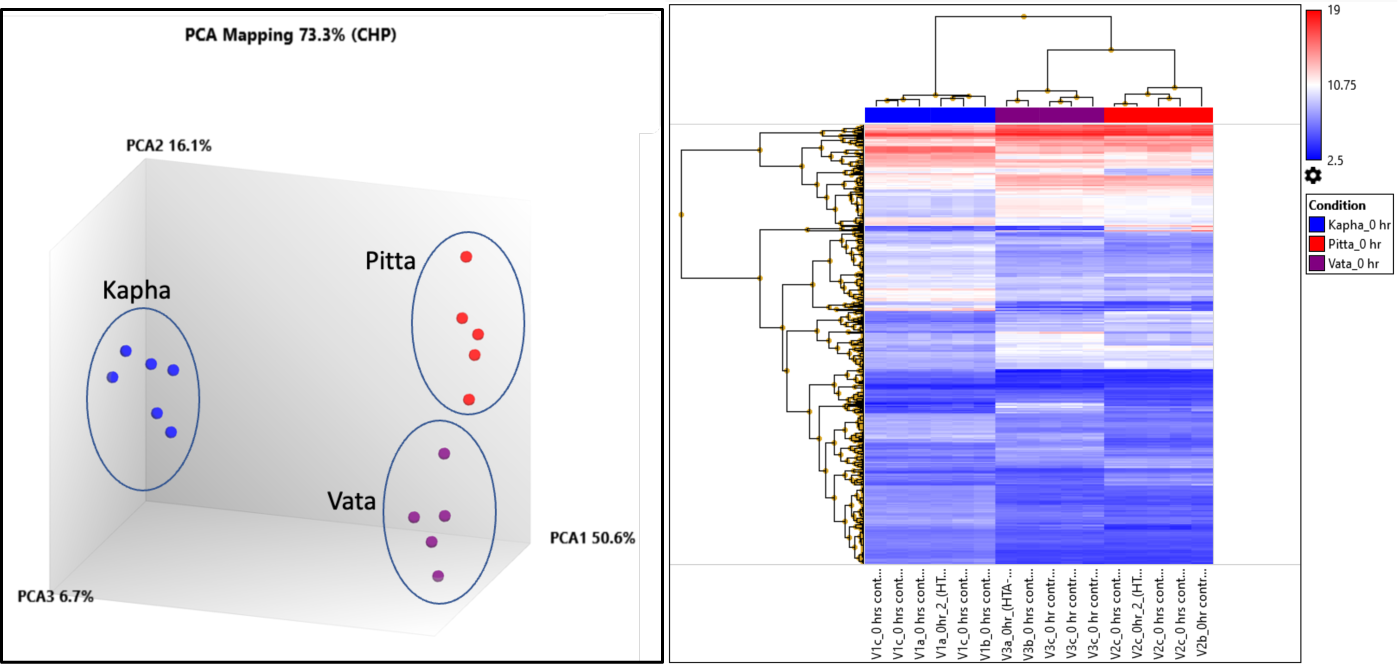
PCA plot and Hierarchical clustering (Condition FDR F-test <0.05) of all 0 hrs. control samples for baseline comparisons among all three *Prakriti* LCLs.

We observed differential enrichment of hallmark gene sets at baseline amongst *Prakriti* comparisons (0 hrs. control samples of all *Prakriti*). Hypoxia as a hallmark gene set was downregulated in Kapha *Prakriti* when compared to Pitta and Vata. Metabolic pathways like glycolysis, heme-metabolism, fatty acid metabolism, and oxidative phosphorylation were also downregulated in Kapha at baseline compared to Pitta and Vata. In Pitta we have observed the higher expression of oxidative phosphorylation both in comparison to Kapha and Vata. The adipogenesis gene set was also downregulated in Kapha as compared to Pitta and Vata.

In Kapha, compared to Pitta and Vata, cell cycle regulators like mitotic spindle and G2M checkpoints were positively enriched at baseline expressions. However, the MTORC1 gene set was downregulated in Kapha when compared to Pitta and Vata. The immune process, like TNF⍺ via NFkB, was negatively enriched in Kapha. The ER stress-induced UPR pathway was downregulated in Kapha compared to Pitta. In contrast, Apoptosis was downregulated in Pitta compared to Kapha and Vata at baseline levels (Fig: 4). Gene enrichment of baseline comparisons using IPA analysis also highlighted differences from immune response to basal metabolic rates among extreme Prakriti groups (Fig: S3 to S5).

**Fig 4:**
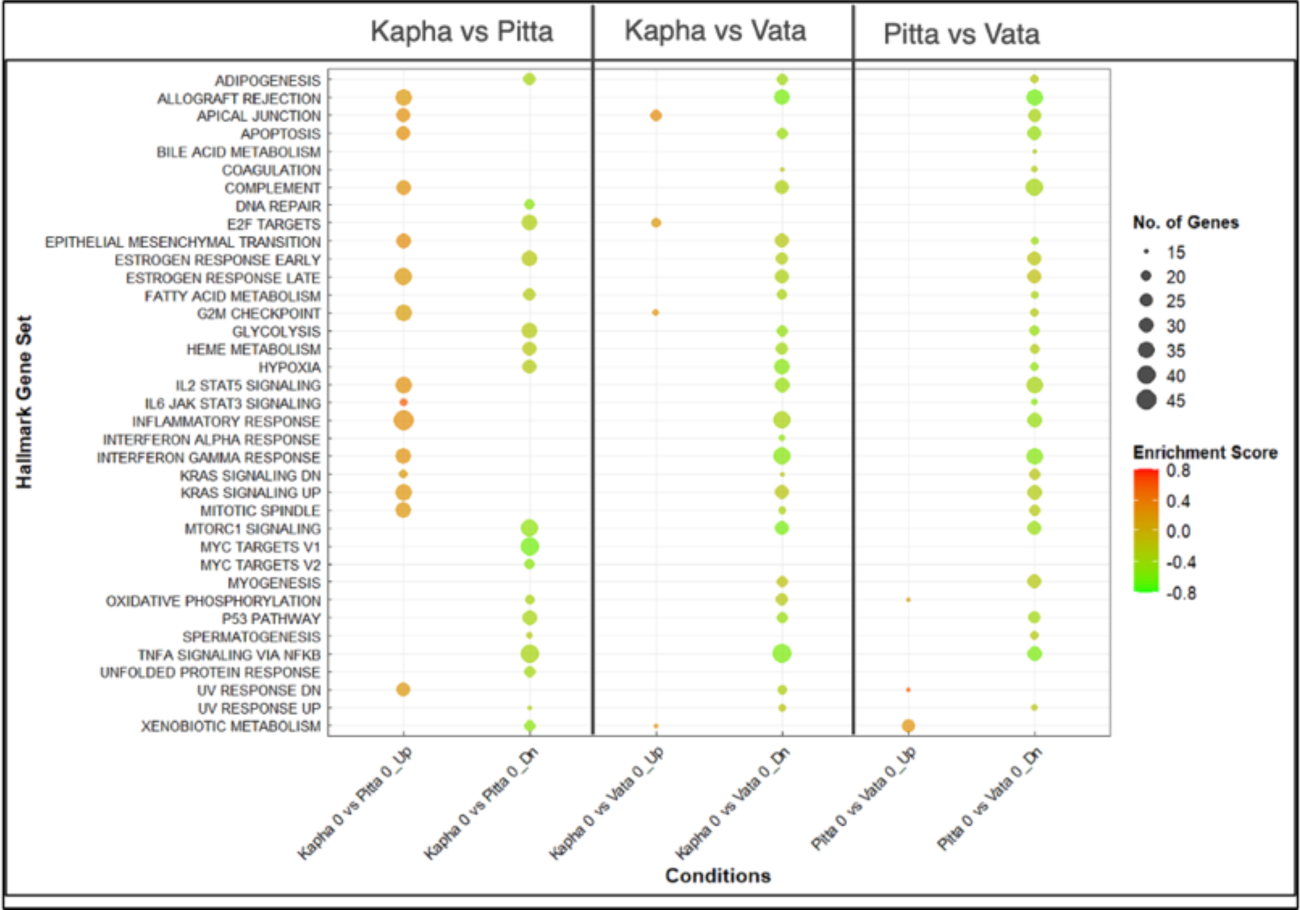
GSEA hallmark gene sets of baseline comparisons of all control 0 hrs. samples in comparison to each other. Nomenclature Kapha 0 vs Pitta 0_Up (Upregulation of hallmark gene set in Kapha as compared to Pitta at 0 hrs), Kapha 0 vs Pitta 0_Dn (Downregulated hallmark gene sets in Kapha as compared to Pitta at 0 hrs.).

### Differential gene expression in response to Hypoxia Pooled data of all LCLs

We compared gene expression patterns of all eight LCLs upon hypoxia treatment at 0.2% oxygen concentration, given for 24 and 48 hrs. and compared with their normoxic time point controls. All hypoxia-treated samples were analyzed in a pooled manner (without *Prakriti* labelling) to determine the overall transcriptomic status of genes at specific time points of hypoxia stress. A total of 5883 and 4776 significant differentially expressed genes (p-value <0.05) were identified after 24 & 48 hrs. of hypoxia. Hypoxia-specific activation of known marker genes, such as *EGLN1, P4HA1, KDM3A, PDK1, VEGFA, SLC2A3, PFKFB3*, and *HK2*, were observed at both time points (supplementary tables). Further, we performed gene enrichment using multiple tools such as Gene Set Enrichment Analysis (GSEA_4.3.1), Ingenuity Pathway Analysis (IPA) and Gene Profiler methods to understand the downstream effects of hypoxia in LCLs in time point specific manner.

Hypoxia-mediated activation of *HIF-1a* signaling was observed at both time points in pooled comparisons (Fig: S8 and S9). In pooled data sets, we observed the overall metabolic reprogramming wherein upregulation of glycolysis, heme-metabolism, cholesterol homeostasis, and downregulation of oxidative phosphorylation and fatty acid metabolism at both time points (Fig: 5 and 6). Activation of the p53 pathway was observed at both time points in pooled data. Gene sets related to the cell cycle and its regulation (MYC, E2F target, G2M checkpoints, cyclin and cell cycle regulator, estrogen-mediated cell cycle entry, and mitotic spindle) and immune response (Allograft rejection, interferon ⍺/ɣ, inflammatory response, and Il-6 JAK STAT3) were down-regulated at both 24 and 48 hrs. time points (Fig: 5 and 6). Gene sets related to nucleotide biosynthesis and DNA repair are also downregulated at both time points in pooled data. Granzyme A is a serine protease that was activated in pooled comparisons after 24 hrs. of hypoxia (Fig: S8). Coagulation is another immune response activated at both time points of continuous hypoxia (Fig: 5 and 6).

**Fig 5:**
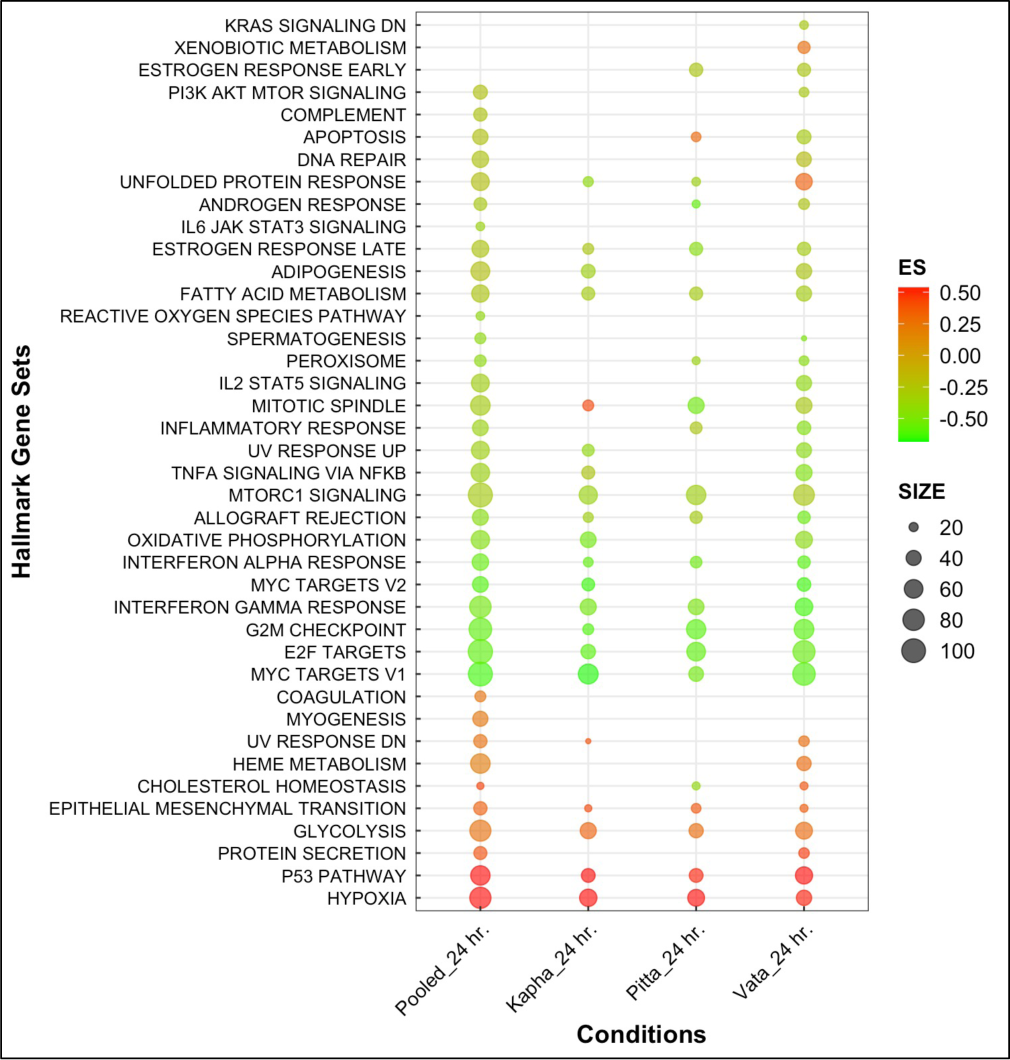
Gene Set Enrichment Analysis of Hallmark gene set for 24 hrs. hypoxia comparisons (p-value<0.05). Pooled (8), Kapha (3), Pitta (2), and Vata (3) analyzed for 48 hrs. time point.

**Fig 6:**
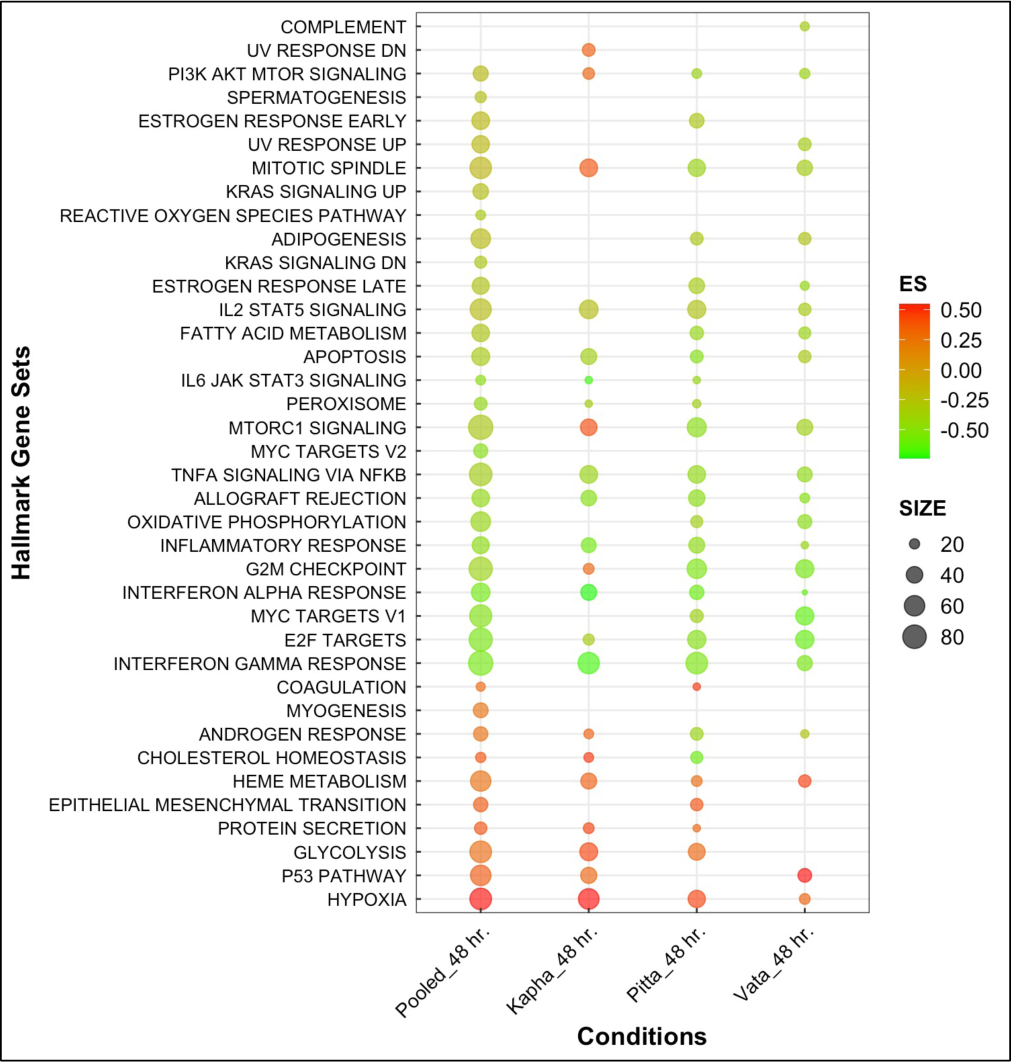
Gene Set Enrichment Analysis of Hallmark gene set for 48 hrs. hypoxia comparisons (p-value<0.05). Pooled (8), Kapha (3), Pitta (2), and Vata (3) analyzed for 48 hrs. time point.

Signaling events related to the activation of immune response like TNF⍺ via NF-kB, IL2-STAT5, and androgen response were downregulated in pooled data sets at both time points (Fig: 5 and 6). The KRAS signaling was specifically down-regulated after 48 hrs. of hypoxia stress in pooled data (Fig: 6). In pooled data, EMT and myogenesis signatures related to developmental stages are activated, while spermatogenesis and adipogenesis are down-regulated at both time points (Fig: 5 and 6). Hypoxia-mediated protective effect on DNA damage response has also been observed after 24 hrs. of stress. Gene set related to the downregulation of UV response was activated, and DNA repair was downregulated after 24 hrs. of hypoxia (Fig: 5).

### *Prakriti*-specific hypoxic response

We further analyzed our data in a *Prakriti*-specific manner to understand the hypoxia responsiveness among individual *Prakriti* groups. DEGs with respect to their normoxic time point controls with significant p-values <0.05 have been used for gene enrichment analysis. In Kapha comparisons, 3506 genes were differentially expressed after 24 hrs. and 3528 genes after 48 hrs. Similarly, in Pitta comparisons, 3440 and 4359 genes were differentially expressed after 24 and 48 hrs. of hypoxia, respectively. In Vata comparison, 4544 genes were differentially expressed after 24 hrs. 3046 genes after 48 hrs. of hypoxia treatment with respect to their normoxic time point controls. These significant differentially expressed genes were further enriched using GSEA and IPA gene enrichment tools. Which have highlighted the major regulatory networks and their respective pathways to understand the *Prakriti*-specific response.

### Hypoxia threshold and extent of fold change

At baseline levels, we observed the negative enrichment of hypoxia hallmark gene sets in the Kapha LCLs compared to the Pitta and Vata LCLs. However, after hypoxia, the maximum positive enrichment was shown in Kapha comparison to their normoxic controls. The extent of fold change and the number of participant genes were higher in the Kapha LCLs. With an enrichment score of 0.53 and 0.54 after 24 and 48 hrs. respectively. Whereas, in Pitta (0.52 and 0.42) and Vata (0.49 and 0.30), enrichment scores were observed for the hypoxia hallmark gene set. At gene levels, for instance, the *KDM3A* gene is among one of the top differentially expressed genes. In Kapha (5.28 and 6.69 FC), in Pitta (3.60 and 3.95 FC), and in Vata (3.56 and 2.84 FC) was observed after 24 and 48 hrs. of hypoxia (supplementary excel).

### Metabolic reprogramming

Hypoxia-mediated metabolic reprogramming was observed in pooled as well as in all three *Prakriti* individually. Hallmark gene sets like glycolysis, gluconeogenesis, were activated as core hypoxic responses at both time points (24 and 48 hrs.) in all comparisons using the GSEA (Fig: 5 and 6). This was also corroborated by enrichment of mitochondrial dysfunction, in canonical pathways of IPA analysis (Fig: S8 to S16). For example, Acetyl CoA biosynthesis (Pyruvate to Acetyl CoA conversion) and mitochondrial oxidative phosphorylation were downregulated at both time points in Vata (Fig: S14 and S15), at 24 hrs. in Kapha (Fig: S10) and at 48 hrs. in Pitta comparisons (Fig: S13). We observed the downregulation of TCA cycle-mediated amino acid regulation. Canonical pathways like methionine degradation in Kapha (Fig S10) and Vata (Fig: S15) and cysteine biosynthesis exclusively in Kapha after 24 hrs. were downregulated (Fig: S10). It would be worth reiterating that there were significant differences between *Prakritis* in metabolic pathways related to the expression gene set. At gene levels, *SLC16A3* (Solute carrier family 16 member 3), one of the monocarboxylate transporter (MCT), transports lactic acid and pyruvate across plasma membrane was varied at baseline comparisons and after hypoxia. we observed the negative enrichment of *SLC16A3* in Kapha compared to Pitta (-1.31 FC) and Vata (-1.2 FC), respectively. However, after hypoxia stress, it was only activated in Kapha (1.39 FC) at 24 hrs. We also observed the activation of heme-metabolism in hypoxic conditions at both time points in the pooled and Vata comparisons, but Kapha and Pitta showed activation after 48 hrs. of hypoxia hallmark gene sets of GSEA (Fig: 6).

### Lipid homeostasis

Pitta LCLs showed downregulation of cholesterol homeostasis and fatty acid metabolism genes at both 24 and 48 hrs. of hypoxia time points as highlighted both in GSEA and IPA analysis (Fig: 5, 6 and Fig: S12, S13). In contrast, the Vata and Kapha groups showed activated cholesterol homeostasis after 24 and 48 hrs., respectively, observed using GSEA analysis (Fig: 5 and 6). However, fatty acid metabolism was downregulated in the Vata LCLs at both time points but in Kapha only after 24 hrs. of hypoxia (Fig: 5 and 6). In Pitta LCLs, downregulation of 3-phosphoinositide biosynthesis and other substrates was observed exclusively after 48 hrs. using canonical pathways of IPA analysis (Fig: S13).

### Cell cycle and death

GSEA highlighted the negative enrichment of cell cycle processes (p53 pathway, Mitotic spindle, G2M checkpoint, E2F target, MYC target genes, etc.) at both 24 and 48 hrs. of hypoxia in pooled comparisons (Fig: 5 and 6). Cell cycle processes like E2F targets were down at both time points, and MYC target genes were down after 24 hrs. in all *Prakriti* comparisons (Fig 5 and 6). However, differences to cell cycle processes are also observed in *Prakriti*-specific manner. In Pitta and Vata LCLs, except the p53 pathway, all cell cycle processes were downregulated at both time points. Whereas, upregulation of mitotic spindle along with p53 pathway was observed in Kapha at both 24 and 48 hrs. of hypoxia. However, G2M checkpoints, which was downregulated after 24 hrs. in Kapha was then upregulated after 48 hrs. of hypoxia. Cell cycle regulators like cyclin and CHK proteins were downregulated after 48 hrs. of hypoxia in Kapha LCLs using canonical pathways of IPA. In contrast, the Vata group LCLs have shown activation of cell cycle checkpoints at 24 hrs., particularly in regulating G1/S checkpoint and p53 signaling to control the progression of the cell cycle (Fig: S14). Kapha and Vata responded differently to hypoxia in terms of cell cycle-related processes. Activation of the GADD45 cell signaling pathway at both time points in Kapha was also observed using IPA analysis (Fig: S10 and S11). Similar observations were also captured using Gene Profiler enrichment of biological processes (Fig: S17 and S18).

### Immune response

Hypoxia-mediated immune signaling processes were also differentially enriched in a *Prakriti*-specific manner. TNF-⍺ via NF-kB signaling was negatively enriched in the Kapha and Vata LCLs after 24 hrs., but it was downregulated in all *Prakriti’s* comparisons after 48 hrs. of hypoxia (Fig: 5 and 6). The IL2-STAT5 signaling was downregulated at both time points in the Vata, but only after 48 hrs. in the Kapha and Pitta LCLs (Fig: 5 and 6). Canonical pathway like PD-1/PD-L1 cancer immunotherapy pathway was activated in Pitta after 48 hrs. using IPA analysis (Fig: S13). Other immune-responsive pathways like TREM1, Th1, IL-12 signaling and production of macrophages are downregulated exclusively after 48 hrs. in Vata comparison through IPA analysis (Fig: S15). We also observed the downregulation of Neutrophil Extracellular Trap (NET), an innate immune response that activates the blood coagulation system was downregulated in Vata LCL lines after 24 hrs (Fig: S14).

Gene Profiler highlighted the downregulation of T-cell receptor and MAPK signaling events were negatively enriched at both time points (24 and 48 hrs.) in Vata LCLs. On the other hand, MAPK was downregulated in Pitta only after 48 hrs. of hypoxia, and no change was observed in the Kapha LCLs. The cytokine-cytokine receptor interaction pathway was downregulated in the Vata at both time points, but other *Prakriti* LCLs showed downregulation only after 48 hrs. of hypoxia. The immune-responsive JAK-STAT signaling pathway showed downregulation after 24 hrs. in the Vata, whereas it was down in the Pitta and Kapha groups after 48 hrs. of hypoxia (Fig: S17 and S18).

### Unfolded Protein Response (UPR)

ER stress-mediated UPR was also regulated in a *Prakriti-*specific manner. GSEA and IPA analysis showed UPR activation in Vata LCLs after 24 and 48 hrs (Fig: 5 and Fig: S15) while Kapha and Pitta observed down-regulation after 24 hrs (Fig: 5) but activated in Kapha after 48 hrs of hypoxia challenge (Fig: S11). Similar observations were also made using the Gene profiler, where activation of UPR was exclusive to Vata after 24 hrs (Fig: S17).

### There were some unique pathways observed to be enriched in one *Prakriti* in response to hypoxia such as follows: TGF-β signaling pathway

Gene enrichment through IPA canonical pathways highlighted hypoxia-mediated activation of the TGF-β signaling exclusively at 24 hrs. in Kapha (Fig: S10). TGF-β induced signaling via SMADs SMAD3 (3.15 FC) and SMAD5 (1.57 FC) was upregulated in Kapha.

### Olfactory transduction

Olfactory transduction was upregulated at both time points in the Pitta and Vata LCLs as observed using Gene Profiler enrichment of biological processes. In contrast, the Kapha group showed downregulation of olfactory transduction after 48 hrs. of hypoxia (Fig: S6 and S7).

### Gene Profiler – Biological processes in *Prakriti*-specific manner

Using Gene Profiler method of gene enrichment of upregulated and downregulated genes separately using fold change >1.5 or < -1.5 with p-value <0.05 significance. In Kapha comparisons, 1772 and 1454 genes were differentially expressed using 1.5-fold change cut off. Similarly, Pitta showed 1325 and 1532 genes and Vata showed 1906 and 1298 genes after 24 and 48 hrs. of hypoxia. This also provided a similar trend as observed in GSEA and IPA, biological processes, such as response to decreased oxygen levels, were upregulated in Kapha and Pitta at both time points after hypoxia. The metabolic reprogramming process i.e., cellular and aerobic respiration, was enriched in downregulated genes of the Vata group after 24 hrs. of hypoxia.

In the Pitta and Vata groups, innate immune processes, including interferon-I, were downregulated after 24 hrs. of hypoxia. Similarly, immune-related processes like regulating hemopoiesis and platelet formation were exclusively downregulated in Vata after 24 hrs. (Fig: S17). Process, like histone lysine demethylation, was upregulated at both time points in Kapha and, after 24 hrs. in Vata cell lines. However, in the Pitta and Vata groups, chromosomal segregation and chromatin remodelling processes were downregulated after 24 hrs. of hypoxia. On the other hand, in the Pitta comparison, processes related to cellular biosynthesis and cellular stress response were downregulated after 24 hrs., and processes related to viral and immune-related processes were downregulated after 48 hrs. of hypoxia. In Gene Profiler data, we also observed the activation of ER to Golgi-mediated vesicle transport through COPII-coated vesicle budding and retrograde transport from Golgi to ER in the Vata group after 24 hrs. of hypoxia similar to GSEA and IPA analysis (Fig: S17).

### qRT-PCR for microarray validation

To validate the microarray data, we performed qRT-PCR of hypoxia-responsive DEG. We have selected the target genes according to their biological significance. For instance, *HIF-1*⍺ and *EGLN1* genes were selected as hypoxia-sensing genes. We have also selected the *HIF1A-AS2* gene, as it was among the top DEGs. It is a lncRNA that can regulate the expression of *HIF-1*⍺ in hypoxic conditions at transcriptomic levels. Similarly, we have selected *SLC2A3*, which is a glucose transporter, *ALDOC*, glycolysis pathway gene, and *PDK1*, which acts as a gatekeeper for the TCA cycle for microarray validation (Fig: 7). We observed a similar expression pattern for all selected genes using qRT-PCR as we observed in microarray data.

**Fig 7:**
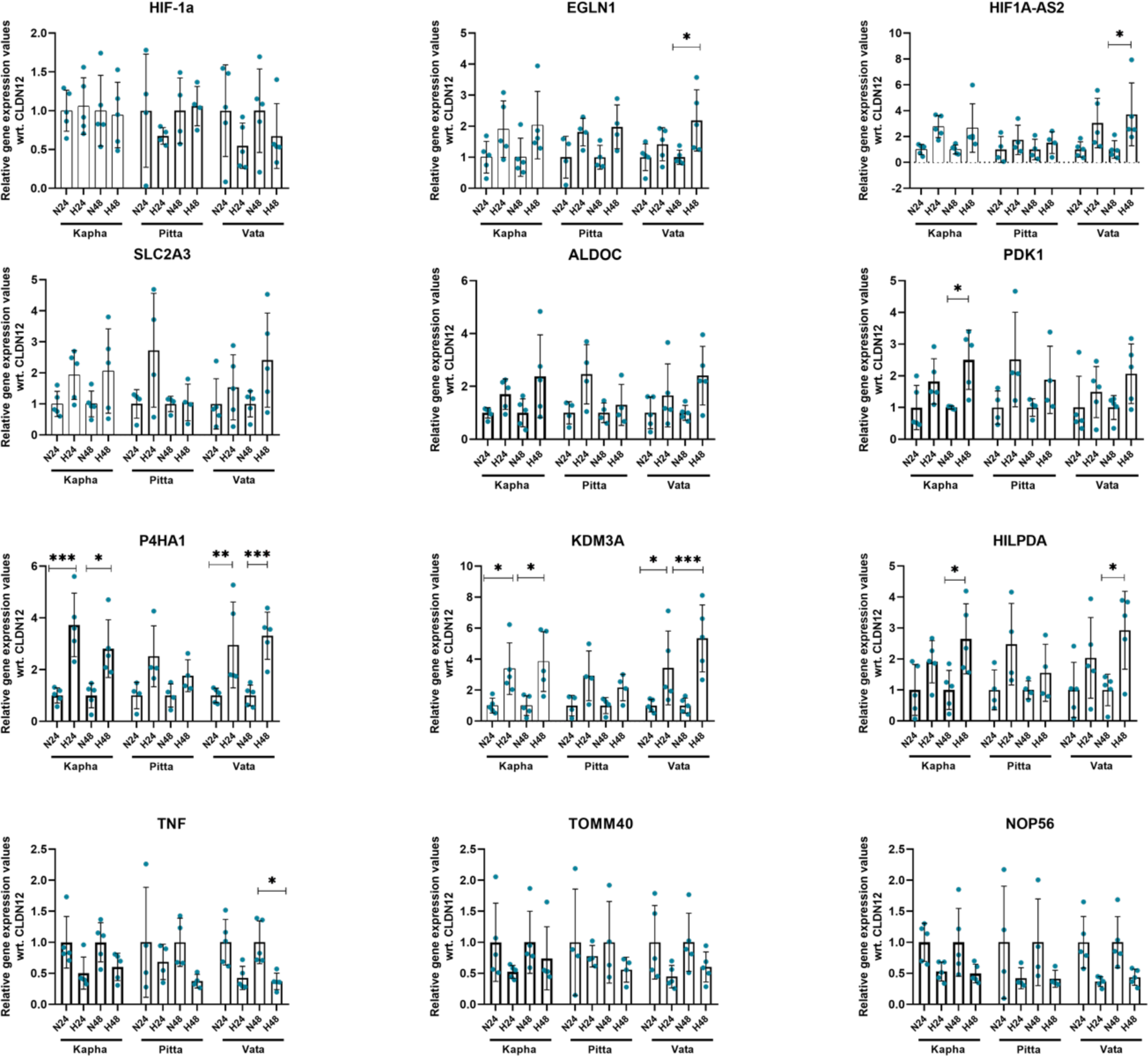
Hypoxia-responsive DEG using the qRT-PCR method in *Prakriti*-specific LCLs (N24 and N48 are normoxic controls, H24 and H48 are hypoxia time points). All eight LCLs are confirmed for hypoxia-specific markers after 24 and 48 hrs. (N=3).

### *Prakriti*-specific cell cycle differences in hypoxic conditions

Gene enrichment of pooled data highlighted the downregulation of overall cell cycle processes, which is also considered a hallmark of energy-deprived conditions. However, we observed the differences in cell cycle processes in a *Prakriti*-specific manner. Kapha LCLs showed activation of cell cycle process in hypoxic conditions along with other hypoxia-specific signatures like higher metabolic reprogramming. In contrast, the Pitta and Vata LCLs showed the downregulation of cell cycle processes at both time points (Fig: 5 and 6). To functionally validate these observations, we tried to capture the cell cycle status using cellular assays in hypoxic conditions at different time points.

### CFSE Assay

CFSE assay is widely used for cell proliferation. It’s a cell staining dye that can dilute over generational shift. We have observed a significant peak shift in Kapha *Prakriti* LCLs after 48 and 72 hrs. in normoxia and hypoxia conditions. In contrast, in Pitta comparisons, peak shifts were not significant and Vata showed a significant peak shift after 72 hrs. of hypoxia. (Fig: 8 and S19).

**Fig 8:**
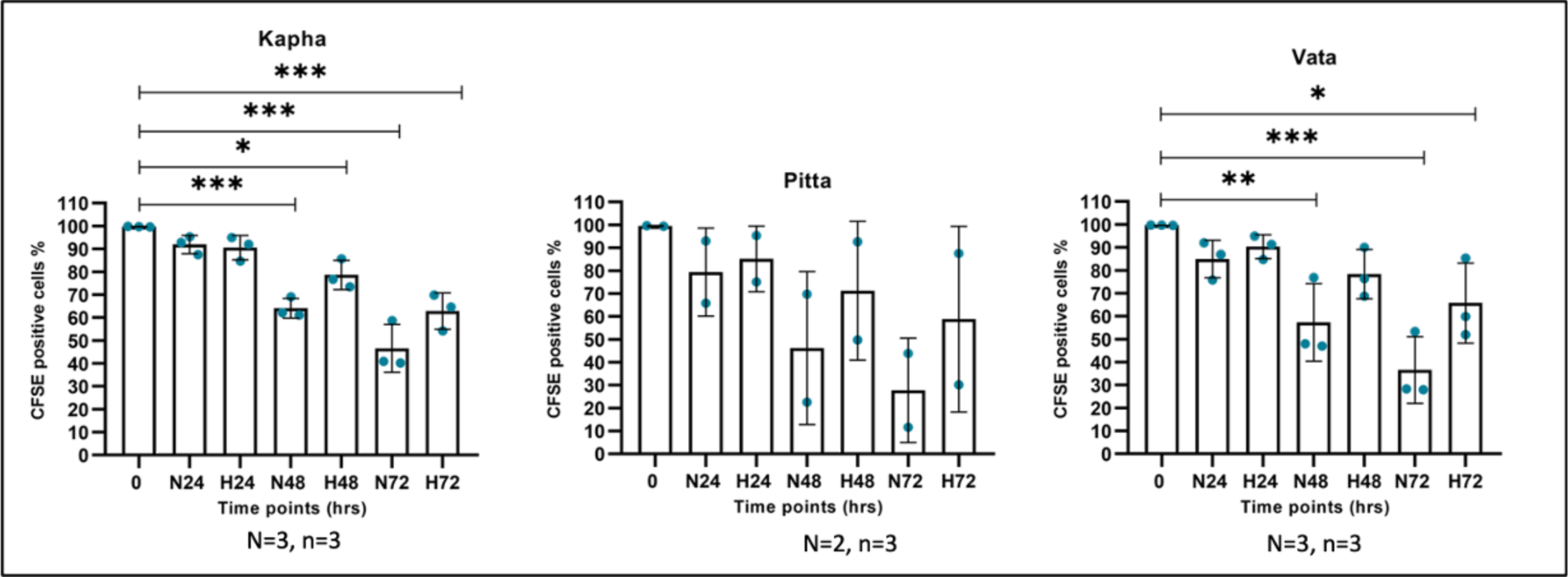
CFSE assay in hypoxic conditions (0.2% oxygen) for different time points (0, 24, 48, and 72 hrs.), where N is normoxic control, and H is hypoxia time points for all eight extreme *Prakriti* LCLs. Hypoxia experiments were performed in three biological replicates followed by acquisition on FACS

## Discussion

In our current work, we studied cellular level variability in hypoxic stress response using the LCLs as a model system derived from healthy individuals of three extreme *Prakriti* types. We have used a total of eight LCLs (3 Kapha, 2 Pitta, and 3 Vata) for hypoxia experiments. *Prakriti*-specific global transcriptomic profile using Affymetrix microarray of Human Transcriptomic Array 2.0 (HTA 2.0) highlighted the overall variability among healthy individuals at baseline and in response to hypoxia. At baseline comparisons, we could identify differences in cellular pathways such as hypoxia and metabolic processes, by comparing untreated (0 hrs. control samples) LCLs in a *Prakriti*-specific manner. Due to baseline differences in these hypoxia-related processes, we considered comparing the hypoxia response of individual *Prakriti* cell lines with their respective normoxic timepoint controls.

Our finding showed that Kapha LCLs have lower expression of hypoxia and p53 pathway hallmark gene sets at baseline comparison. In addition, the lower expression of metabolic processes was observed in Kapha compared to Pitta and Vata at baseline levels. At baseline levels, gene sets related to metabolic reprogramming, such as glycolysis, oxidative phosphorylation, fatty acid metabolism and adipogenesis, have lower expression in Kapha as compared to 0 hrs. controls of Pitta and Vata. In contrast, it showed maximum change in metabolic processes after hypoxia when we compared *Prakriti*-specific LCL lines with their normoxic time-point for 24 and 48 hrs. using 0.2% oxygen concentrations. For example, negative enrichment of metabolic pathways like oxidative phosphorylation, fatty acid metabolism, and adipogenesis were observed in Kapha after 24 hrs. of hypoxia. However, Pitta and Vata have downregulation to these energy-consuming metabolic processes after 48 hrs. of hypoxia. Oxidative phosphorylation, which was higher in Pitta at baseline (*CYB5A, CPT1A, NDUFS6*), was downregulated after 48 hrs. of hypoxia, whereas in Kapha, it had lower expression in comparison to the other two at baseline and responded to hypoxia by downregulating it further after 24 hrs. TCA cycle-mediated amino acid regulation was also downregulated exclusively in Kapha LCLs. Glycolysis was activated in Vata only at 24 hrs., but others have upregulation at both time points. Hypoxia response was activated in all 3 *Prakritis*, although HIF-1⍺ expression was not observed in any of the cell lines, similar to other chronic hypoxia studies at different cell types(15) at selected time points. However, we observed the significant activation of hypoxia hallmark gene sets upon hypoxia in all three *Prakritis* at 24 and 48 hrs.

Lipid homeostasis is crucial to promote cell survival response in hypoxic conditions. It is known to be altered in hypoxic conditions to promote an anaerobic mode of respiration to rescue the energy-deprived conditions(16). Cholesterol homeostasis is an energy-dependent process that plays a vital role in regulating membrane fluidity and is downregulated in Pitta at both time points. Hypoxia-mediated degradation of SREBP2, which is an initiating factor for cholesterol synthesis, protects cell death and forces cancer cells to take up exogenous cholesterol instead of synthesizing it(17). Another membrane phospholipid, 3-phosphoinositide, was also downregulated in Pitta at 48 hrs. of hypoxia. Phosphoinositides constitute a network of soluble and membrane-associated inositol phosphates. They play a crucial role in coordinating cellular responses, from nutrient uptake and utilization to growth factor signaling and energy homeostasis(18). Alteration of lipid homeostasis in hypoxic conditions is required to cope with energy-deprived conditions, but this can also affect membrane fluidity, resulting in leaky channels, which can further activate downstream immune response. Altered lipid homeostasis was exclusively downregulated in Pitta comparisons.

Pitta exhibited lower gene expression levels for immunological processes, while Vata showed higher levels at baseline levels. After hypoxia, all three *Prakritis* demonstrated downregulation of immune response, but in a time-specific manner. For example, TNF⍺ via NFkB, which was initially lower in Kapha, was further downregulated in response to hypoxia at 24 and 48 hrs. In contrast, Pitta, which has higher baseline levels than Kapha, showed negative enrichment only after 48 hrs. of hypoxia. In Vata LCLs, higher expression of immunological processes at baseline showed negative enrichment to immunological processes at both time points. However, number of participating genes were low at 48 hrs. in Vata for immunological processes as compared to their normoxic controls. Hypoxia triggers the MAPK pathway by activating ERK kinases through phosphorylation and nuclear translocation(19). However, we have observed the downregulation of the MAPK pathway in the Pitta and Vata groups. The MAPK protein kinase family are crucial in various cellular processes such as cell growth, survival, differentiation, development, cell cycle regulation, and cell death. We have observed the downregulation of UVB-induced MAPK signaling in Vata LCLs after 24 hrs. Similarly, MAPK-activated ERK5 (Extracellular signal-regulated kinase 5) induces a variety of stimuli such as growth factors, GPCRs and cellular stress factors like oxidative stress. Hypoxia also suppresses the production of type I IFN but not nuclear factor-κB–dependent proinflammatory cytokines(20). We observed the downregulation of inflammatory alpha and gamma response in all comparisons in pooled as well in a *Prakriti*-specific manner. However, their number of participating genes were different from one comparison to another. This temporal aspect of the immune response provides valuable insights into the dynamics of the immune system in different *Prakritis*.

Similar to our previous published study(14), we have observed baseline differences in cell cycle-related processes among extreme *Prakriti*-derived LCLs. We observed expression level differences in cell cycle processes in Kapha LCLs compared to Pitta. Processes like G2M checkpoints, Mitotic spindle, and KRAS signaling were positively enriched, and processes like E2F and MYC target gene sets were negatively enriched in Kapha compared to Pitta at baseline. However, in Vata comparisons, baseline cell cycle-related processes were upregulated as compared to other *Prakritis*. In contrast, after hypoxia, we observed the overall downregulation of cell cycle processes at both time points, in all LCLs except for Kapha. Cell cycle processes like, G2M checkpoints and mitotic spindle are activated in Kapha as compared to their normoxic controls. Similarly, cell proliferation specific, KRAS and MYC were downregulated in Vata at selected time points. Cell cycle processes and their checkpoints, which are known to be suppressed in hypoxic conditions(21), were still activated in Kapha comparisons at both time points. In the Pitta and Vata groups, except for the p53 pathway, all other cell cycle-related processes were downregulated at both time points. The p53 gene is considered to be the central mediator of the DNA damage response. Under hypoxic conditions, p53 stabilizes and activates the GADD45 (Growth Arrest and DNA Damage-inducible 45) signaling pathway. Activated GADD45 signaling promotes cell survival by inducing cell cycle arrest and promoting DNA repair through a variety of stress and growth regulatory mechanisms, including G2M checkpoint control. However, in persistent stress conditions, GADD45 initiate apoptosis by initiating JNK/p38 MAPK signaling(22). Activation of the GADD45 cell signaling pathway was observed exclusively at both time points in Kapha LCLs.

Hypoxia and other cellular stressors can lead to extensive protein modification, which accumulates unfolded or misfolded proteins in the ER lumen. Clearance and regulation of the unfolded proteins are crucial to promote cell survival responses(23). Activation of the UPR was observed at both time points in the Vata group and in Kapha after 48 hrs. of hypoxia. Genes like *ASNS, ATF3,* and *ATF4* from PERK-mediated UPR pathways are activated in Vata. However, at baseline, UPR gene set expression was higher in Pitta than in Kapha, but no change was observed in Pitta after hypoxia induction. Regulation of the UPR to hypoxia highlights the ongoing rescue mechanisms in different groups. UPR activation will lead to the activation of cell survival mechanisms through autophagy; if the condition persists, it will also induce cell death(24). The PI3K-mTOR signaling regulates the upstream activity of HIF-⍺ by promoting its mRNA expression. In addition, it also regulates the expression of angiogenic factors in cancer cells via the HIF-1⍺ axis-mediated downstream activation of the target genes(25). We observed the activation of PI3K-AKT-mTOR signaling in Kapha after 48 hrs. of hypoxia; in contrast to other groups, including pooled comparison, it was downregulated after 48 hrs. of hypoxia.

Cells evoke survival responses by promoting alternative and efficient ways of cellular energy utilization during hypoxic conditions. Activating TGF-β signaling is one such example. It was activated exclusively after 24 hrs. in Kapha, which can further promote downstream signaling by directly inducing ROS levels to promote HIF-⍺ stabilization and indirectly by fostering glycolysis in tumour cells through SMAD3-mediated pathway(26). TGF-β signaling also promotes profibrotic processes by activating ⍺-SMA levels directly or HIF-dependent manner. Studies on cell lines have demonstrated that HIF-1α/TGF-β activation under hypoxic conditions leads to increased levels of α-SMA and other myofibroblast markers. Studies have also highlighted the role of anaerobic respiration as a critical factor in myofibroblast differentiation in pulmonary fibrosis(27). Hypoxia-mediated induction of TGF-β in relevant cellular models could further help to understand its role in disease progression and severity. Another pathway is olfactory transduction. Olfactory receptors are highly expressed in carotid bodies, which are the primary sensor of arterial blood oxygen concentrations(28). We observed the activation of olfactory transduction in *Prakriti*-specific manner. Kapha LCLs have downregulated the olfactory transduction pathway at 48 hrs., whereas, in Pitta and Vata, it was upregulated at both time points.

We have observed the hypoxia-mediated activation of lncRNA HIF1A-AS2 is required for cell proliferation, epithelial-mesenchymal transition (EMT) and tumour propagation in non-small cell lung cancer (NSCLC). Transcriptomic analysis of *HIF1A-AS2* reveals its role as a trans modulator of gene expression, particularly regulating transcriptional factor genes, including MYC. It activates MYC by recruiting DHX9 on its promoter, consequently stimulating the transcription of MYC and its target genes(29). We observed the higher expression of HIF1A-AS2 lnc RNA at both time points in Kapha up to 6 fold from their respective time point control. In Pitta and Vata, upto 3-fold was observed after 24 hrs. Downregulation of the MYC (-2.27 fold) gene was exclusively observed in Pitta after 48 hrs. only.

We have observed that the Kapha group showed consistent activation and regulation of cell cycle processes from baseline levels to energy-deprived hypoxic conditions. To functionally validate these observations, we used a CFSE cell proliferation assay. Cell proliferation rates were similar after 24 hrs. in all groups after hypoxia as compared to their time-point control. However, in the Kapha group, we have seen a significant peak shift after 48 and 72 hrs. in both normoxic and hypoxic conditions. In contrast, in the Pitta and Vata group, hypoxia samples showed low peak shift as compared to their time-point control. So, this also corroborates with our previous observations that Kapha had maintained the cell proliferation machinery even in hypoxic conditions.

## Conclusion

The cellular response to hypoxia was studied using *Prakriti*-specific cell lines. Our observations highlighted that while the core response to hypoxia is observed in all individual cell lines, there are specific differences in the way each *Prakriti* responds and the trajectory it could take at the cellular levels. Differences in the impact on pathways and processes are apparent, along with unique responses. This could be due to differences in their baseline sensitivity and metabolic patterns, as evidenced by the baseline differences in some core hypoxia and metabolism-related genes. In the current study, Kapha *Prakriti*, which had lower expression of metabolic process and hypoxia-responsive genes at baseline, showed maximum change in hypoxia-mediated metabolic reprogramming at both time points. We observed differences in the number of participating genes and their existing fold change compared to other *Prakritis* to metabolic processes. Despite facing energy deprivation, Kapha regulated their cell proliferation in hypoxic conditions, whereas others have been completely downregulated at both time points. The activation of the TGF-β signaling and PI3K/AKT/mTOR pathway in Kapha highlighted the alternative approach to cell survival response to hypoxia. On the other hand, in Pitta *Prakriti*, along with activation of the core hypoxia response, downregulation of cholesterol homeostasis was an alternative cell survival response. On the other hand, Kapha and Vata have activation of cholesterol homeostasis in a time point specific manner. The downregulation of Phosphoinositide exclusively in the Pitta comparison also highlighted its effect on membrane liquidity and integrity in hypoxic conditions. In Vata *Prakriti*, higher expression of immune-responsive genes at baseline levels experienced maximum downregulation after hypoxia at both time points. Vata *Prakriti* LCLs have shown an alternative response mechanism by activating the UPR pathway at both time points of hypoxia.

Our work towards understanding hypoxia sensing and response among healthy individuals of extreme *Prakriti* types will provide a fresh perspective toward defining new baseline levels and exploring inter-individual variability among hypoxia-driven disease conditions. To explore the disease mechanism and its molecular targets among diverse populations with differential outcomes, one can utilize the non-invasive multi-phenotyping-based *Prakriti*-stratification method of Ayurveda. Functional validation of hypoxia outcomes in different cell types in a *Prakriti*-specific manner would be important to explain the inter-individual response to hypoxia and predict its outcomes.

## Methodology

### *Prakriti* specific Lymphoblastoid cellular models

Our current study used previously developed extreme *Prakriti*-derived LCLs in our lab to explore the hypoxia sensing and response pathways. A total of eight LCLs were developed using EBV transfection of PBMCs and characterized at morphological, genetic, and cellular levels. LCLs were cultured using RPMI (Roswell Park Media Institute) media with 15% Fetal Bovine Serum (FBS) and antibiotics (1X Anti-Anti) reagents. Cellular morphology and growth have been observed during the culturing and maintenance of LCLs. Cell sterility has been routinely confirmed for mycoplasma contamination using a mycoplasma detection kit (Lonza, cat #: LT07-318).

### Hypoxia generation and time point validation

The exact oxygen concentrations for the induction of hypoxia response in any cell type depend on several factors such as cell type, metabolic rate, energy demand and duration of hypoxia etc. So, it is crucial to ensure hypoxia generation in the cellular models to evoke a hypoxia response. Chronic hypoxia at cellular levels is required to study expression level differences. We have not come across any such studies where LCLs have been used for hypoxia-specific transcriptomic signatures, so we looked for other cellular studies. Through literature search, we have encountered different cellular studies using 0.2% oxygen concentration at different time points to develop chronic hypoxia (from 16 hrs. and above) at cellular levels (30,31). We used chamber-mediated hypoxia with a 0.2% oxygen concentration in LCLs models. LCLs were screened at different time points from 0 to 72 hrs. (0, 2, 4, 6, 12, 24, 48, and 72 hrs.) at 0.2 % oxygen concentration, followed by the detection of hypoxia markers using metabolic reprogramming in pooled and qRT-PCR.

### Western Blotting

For western blotting experiments, LCLs were seeded (1×10^6^ cells) in untreated 6-well culture plates in duplicates for 0 to 72 hrs. time points. One plate was kept in normoxic conditions at 21% oxygen and another in the hypoxia chamber at 0.2% oxygen concentration, and protein lysate was prepared using RIPA (Radioimmunoprecipitation Assay) (Sigma, cat no. R0278) buffer along with a Protease inhibitor cocktail (PIC) (Sigma, cat no. P8340) and Dithiothreitol (DTT) in 1000:10:10 ratio. After treatment, all cellular handling protein lysate preparation was performed at 4°C to prevent cellular hypoxic conditions. Once lysate was prepared, it was quantified using a colourimetry assay called BCA (Bicinchoninic Acid) protein estimation kit from Thermo (Cat No. 23225) using a spectrophotometer.

For hypoxia-specific target protein (HIF-1⍺ and EGLN1) detection in western blotting, we have used 60 µg of total protein concentration, along with a protein ladder (Thermo cat no. 26619, 26616), loaded with over 12 % SDS PAGE gel. Proteins were transferred to the PVDF membrane (MDPI, cat no. SVFX8301XXXX101) at 30V for overnight transfer at 4℃ using. We have used 5% skimmed milk (Himedia, cat no. GRM1254) for blocking and antibody dilutions. Blocking has been performed at room temperature for 2 hrs., followed by primary antibody treatments overnight at 4 degrees Celsius. Hypoxia-specific antibodies for HIF-1⍺ (Abcam, cat no. ab51608) and PHD2 (Abcam, ab4561) (1:1000 dilution) have been used along with β-actin (Sigma, cat no. A5441) (1:5000) as a housekeeping protein. Blots were developed using Enhanced Chemiluminescence (ECL) reagent (Abcam, cat no. ab133406). Western blotting data were analyzed using Image J software.

### qRT-PCR

We have used the qRT-PCR method to find the expression pattern of HIF-1a and EGLN1 markers for the given time points 0, 2, 4, 6, 12, 24, 48, and 72 hrs. We have used 0.2% oxygen concentration for the hypoxia dose. LCLs were seeded (1×10^6^ cells) in duplicates in untreated 6-well plates for the hypoxia experiment. One plate is kept in normoxia, and the other is kept in the hypoxia chamber. It is known that hypoxia will revert within a few minutes once it comes in normoxic conditions. So, to prevent hypoxia-induced transcriptional changes, cells were immediately transferred to ice. Total RNA has been isolated using the TRIzol method with chloroform and isopropanol. Samples were checked for their integrity and concentration using 1% agarose gel and nanodrop. Further, RNA samples have been treated with DNAse to remove any DNA contaminations using Turbo DNA free kit from Thermo, cat no. AM1907. cDNA synthesis was performed using a High-capacity cDNA synthesis kit (Thermo, cat no.4368814). Primers were designed to capture hypoxia activation at transcriptional levels. We have selected two genes, hypoxia-responsive *HIF-1⍺* and master oxygen sensor *EGLN1,* for transcription level confirmation. Transcriptional sequences of the selected genes were obtained from the Ensembl genome browser. All primers were designed using the NCBI primer designing tool, Primer 3 (version) (Table: 1). Primer specificity has also been confirmed using MFE primer 3.0 software. Primer specificity has been confirmed at different cDNA concentrations.

**Table 1:**
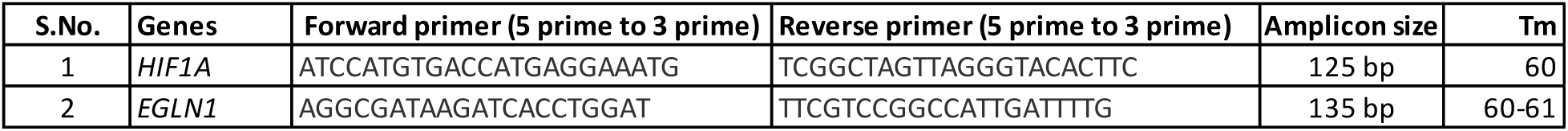
Primer sequences for hypoxia generation and validation using qRT-PCR.

### Gene expression (microarray) at selected time points

We have used the Affymetrix microarray platform to study the global gene expression patterns in response to hypoxia. Untreated LCLs at 0 hrs. for baseline comparisons and hypoxia 24 (H24) and 48 hrs (H48) time points were selected for microarray along with their normoxic experimental controls (N24 and N48). LCLs were seeded in untreated 6-well plates with 1.5 × 10^6^ cells for the experiments. On the day of the experiment, we confirmed the cell number and their health through microscopy. Cells were harvested for RNA isolation after hypoxia treatment (24 and 48 hrs.). We immediately transferred cells to ice for hypoxia time points to prevent the ongoing molecular processes. Further RNA isolation has been done using the TRIzol method, and RNA integrity and concentration have been confirmed through 1% Agarose gel and Nanodrop quantification. A Human Transcriptomic Array 2.0 (HTA 2.0) chip (Affymetrix, cat no. 902162) from the Affymetrix microarray platform has been used for gene expression profiling. For the microarray, 250 ng of total RNA was used as per the manufacturer’s protocol. After ssDNA preparation, it hybridised to the chip for 16 hrs. at 45 degrees in a shaking incubator. The chips were washed and stained using the Affymetrix fluidics station. The chips were scanned, and CEL files were generated using an Affymetrix scanner.

### Transcriptomic analysis

Gene expression analysis was performed using Affymetrix Transcription Analysis Console (TAC) software. CEL files were used as input data, normalization and background correction were done using the gene-level SST-RMA normalization method. The CHP files were generated, which were compared to get the differentially expressed genes. As we have performed our microarray in different batches, we have performed Exploratory Group Analysis (EGA) for batch effect removal using TAC software. Differentially expressed genes were analyzed using CHP files after batch effect removal. For our further analysis, we applied two approaches; in the first one, we pooled all data in terms of hypoxia time points manner to explore the overall transcriptomic differences in response to hypoxia at 24 and 48 hrs. In the second approach, we analyzed it in a *Prakriti*-specific manner, where we observed the hypoxia response within *Prakriti* groups at 24 and 48 hrs. compared to its normoxic controls.

### Gene ontology-based analysis of differentially expressed genes

We have used multiple tools for gene ontological enrichment analysis of differentially expressed genes. For instance, in Gene profiler-based analysis, we took genes with a p-value < 0.05 and fold change cut-off value of 1.5 fold and above or -1.5 fold or below for biological processes and pathways enrichment(32). Secondly, we used genes with a significant p-value cut-off value (p-value < 0.05) for gene enrichment through Gene Set Enrichment Analysis (GSEA)(33) and Ingenuity Pathway Analysis (IPA)(34) software. In GSEA, enrichment of significant differentially expressed genes was performed using ranked gene lists arranged based on their fold change differences from highest to lowest. Hallmark gene sets have been used to enrich hypoxia-responsive pathways, which are the curated gene sets from the 33196 gene sets of the molecular signature database of GSEA. We performed IPA-based enrichment analysis for canonical pathways, which are updated databases for metabolic and cell signaling pathways.

### Microarray validation using qRT-PCR

Affymetrix expression data of differentially expressed genes has been validated using qRT-PCR. We have used the qRT-PCR method to validate the hypoxia-responsive differentially expressed genes. We used 0.2% oxygen concentration for the hypoxia dose. LCLs were seeded in duplicates in untreated 6-well plates for selected hypoxia time points of 24 and 48 hrs. of hypoxia. One plate is kept in normoxia, and the other is kept in the hypoxia chamber. After hypoxia treatment, cells were immediately transferred to ice and proceeded for RNA isolation. Total RNA has been isolated using the TRIzol method with chloroform and isopropanol. Samples were checked for integrity and concentration using 1% agarose gel and nanodrop. Further, RNA samples were treated with DNAse to remove any DNA contaminations using a Turbo DNA-free kit from Thermo, cat no. AM1907. cDNA synthesis was performed using a High-capacity cDNA synthesis kit (Thermo, cat no.4368814). Primers were designed to capture hypoxia activation at transcriptional levels. A total of 12 genes were selected for qRT-PCR validation (Table no. 2). Hypoxia-specific markers, such as *EGLN1* for hypoxia sensing and *HIF-1⍺* for transcriptional activation of hypoxia, have been selected. Other markers, such as *SLC2A3* and *PDK1*, representing regulators of metabolic reprogramming, were selected to confirm hypoxia generation at given time points (24 and 48 hrs.). Transcriptional sequences of the selected genes were obtained from the Ensembl genome browser. All primers were designed using the NCBI primer designing tool, Primer 3 (version). Primer specificity has also been confirmed using MFE primer 3.0 software. Primer specificity has been confirmed at different cDNA concentrations.

**Table 2:**
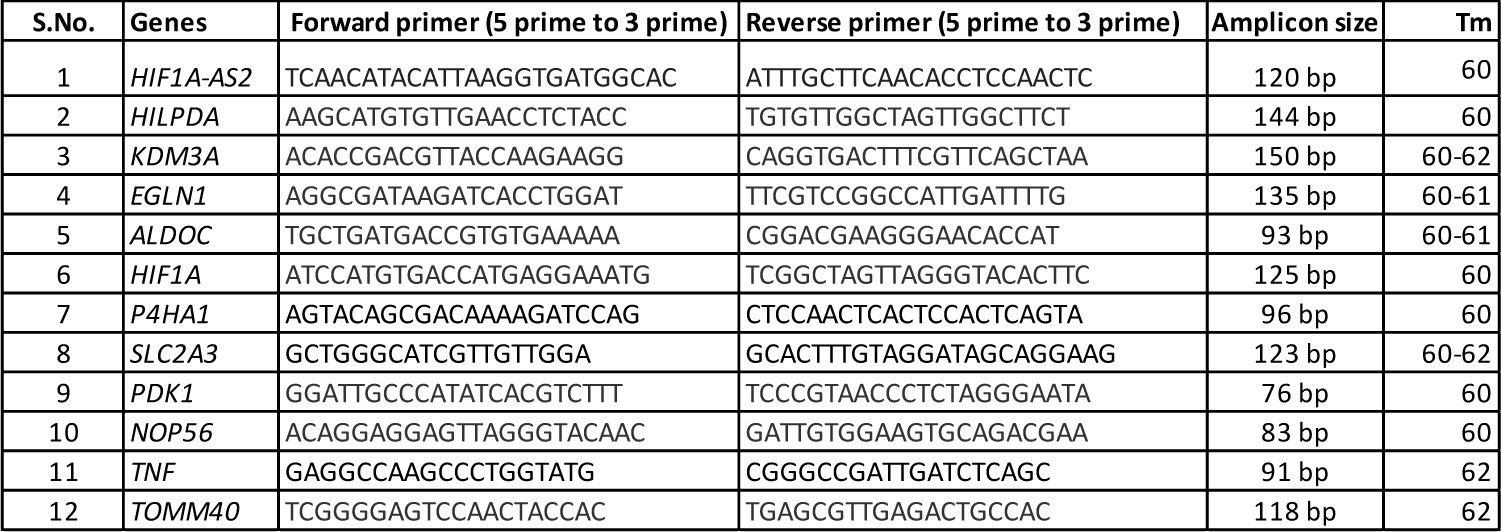
Genes and their primer sequence for microarray validation using qRT-PCR.

### CFSE (Carboxyfluorescein succinimidyl ester) Assay

CFSE is a commonly used cell assay to detect the cell proliferation status among cell lines (Thermo cat no: C34554). It is a dye that can bind to the cells and dilute over generational shifts. Cell media was changed one day prior to ensure proper cell proliferation and growth. On experiment day, cells were treated with 2.5 µM CFSE dye to the cells in 1X PBS, followed by 20 mins incubation at a CO^2^ Incubator. Next, we washed the cells with 1X PBS to remove excess dye. Cells were seeded in fresh RPMI media and kept it in a CO^2^ incubator for 30 mins. before hypoxia treatment. In the next phase, cells were placed in hypoxia incubator along with their time-point controls. The 0 hrs. sample was harvested during the hypoxia incubation as a control sample, where CFSE-positive cells were measured using the FACS method. Cells were given hypoxia treatment from 24 to 72 hrs. and compared with their normoxic controls. Peak shifts were measured through the FACS.

**Supplementary data**: https://doi.org/10.5281/zenodo.11618704 Microarray raw data (CEL files) has been submitted to GEO (GSE235776).

## Author Contributions

BP conceived and designed the study. DS and SKA designed the initial hypoxia experiments. DS performed hypoxia and microarray experiments followed by gene enrichment analysis. DS and KM performed the functional validation experiments. RR prepared the graphs of microarray gene enrichment data. DS and BP wrote the manuscript. BP reviewed the data and manuscript. All authors have read and approved the manuscript.

## Supporting information

10.5281/zenodo.13136722

## Acknowledgements

We express our gratitude to Dr. Mitali Mukerji for her guidance during the conceiving and designing of the study. We are thankful to Dr. Rituja Patil and Dr. Sunjay Juvekar for their contribution to volunteer recruitment. We are thankful to Dr. Sunanda Singhmar and Dr. Sumita Chakarborty for their assistance during LCL culture and valuable input during initial hypoxia experiments. Our thanks also go to Dr. M. Faruq and Pooja for their support during Affymetrix microarray experiments. Additionally, we appreciate the contributions of Dr. Rajesh Pandey and Dr. Ishaan Gupta for their assistance in microarray data analysis and validation experiments. Special thanks to Dr. Soumya Sinha Roy, Dr. Sivaprakash Ramalingam, and Dr. Sangam Giri Goswami for their insightful suggestions during functional validation experiments. We are also grateful to Aditya Iyer and Pragya Gupta for their assistance during the functional validation of hypoxia leads through FACS. Lastly, we would like to extend our sincere appreciation to Dr. Charu Lata and Dr. Benazir Chisti for their invaluable input during manuscript writing and final draft preparation. DS and RR acknowledge the Indian Council of Medical Research (ICMR) for SRF fellowship.

## Ethics Statement

The protocol was approved by the Institutional Human Ethics Committee (IHEC) and Institutional Biosafety committee (IBSC) of CSIR-Institute of Genomics and Integrative Biology (IGIB), New Delhi, India.

