## Supplementary material for "Transcriptomic profiling of healthy individual-derived LCLs revealed inter-individual variability towards-hypoxia-responsive pathways": 10.5281/zenodo.13136722

#### Supplementary data

Table S1: Summary of eight Prakriti-specific Lymphoblastoid cell lines (LCLs)

| S.No. | Prakriti | Gender | Age | LCL Code |
|---|---|---|---|---|
| 1 | Kapha | Male | 30 | LCL V1a |
| 2 | Kapha | Male | 28 | LCL V1b |
| 3 | Kapha | Male | 29 | LCL V1c |
| 4 | Pitta | Male | 23 | LCL V2b |
| 5 | Pitta | Male | 37 | LCL V2c |
| 6 | Vata | Male | 28 | LCL V3a |
| 7 | Vata | Male | 28 | LCL V3b |
| 8 | Vata | Male | 35 | LCL V3c |

#### Hypoxia generation and time point validation

Fig S1(a)

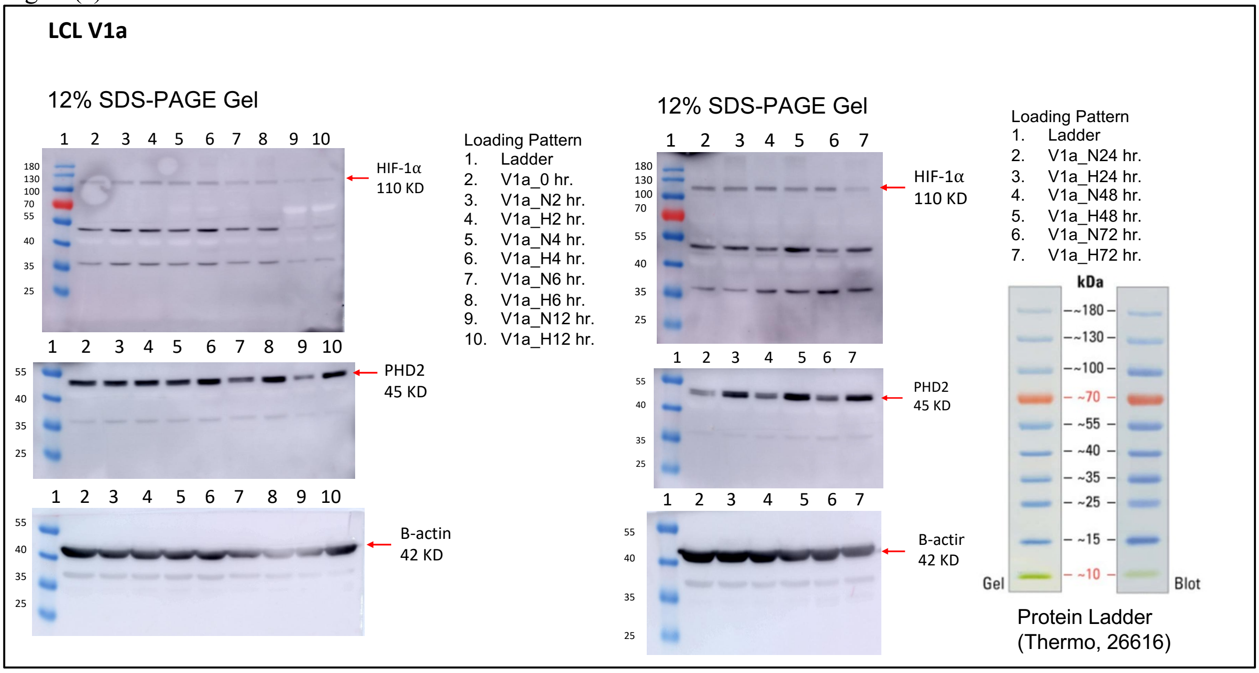

Fig S1(b)

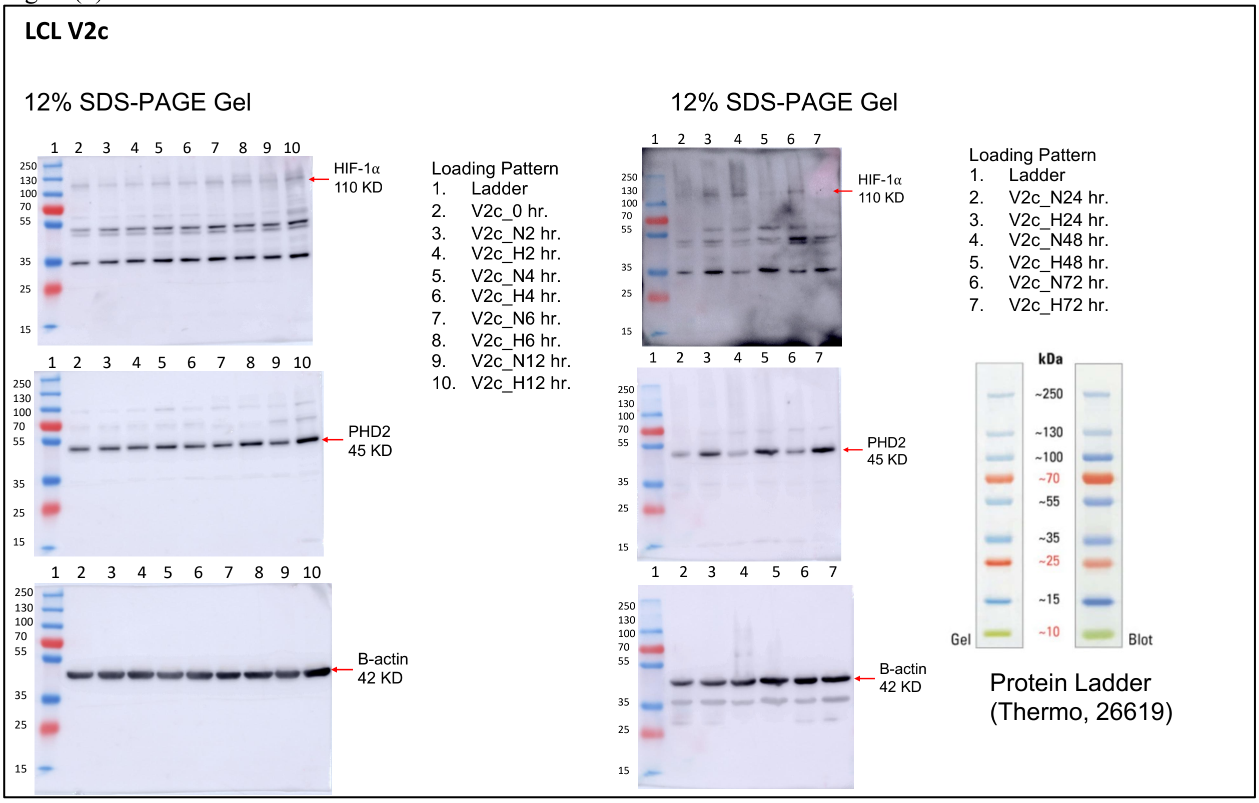


Fig S1(c)

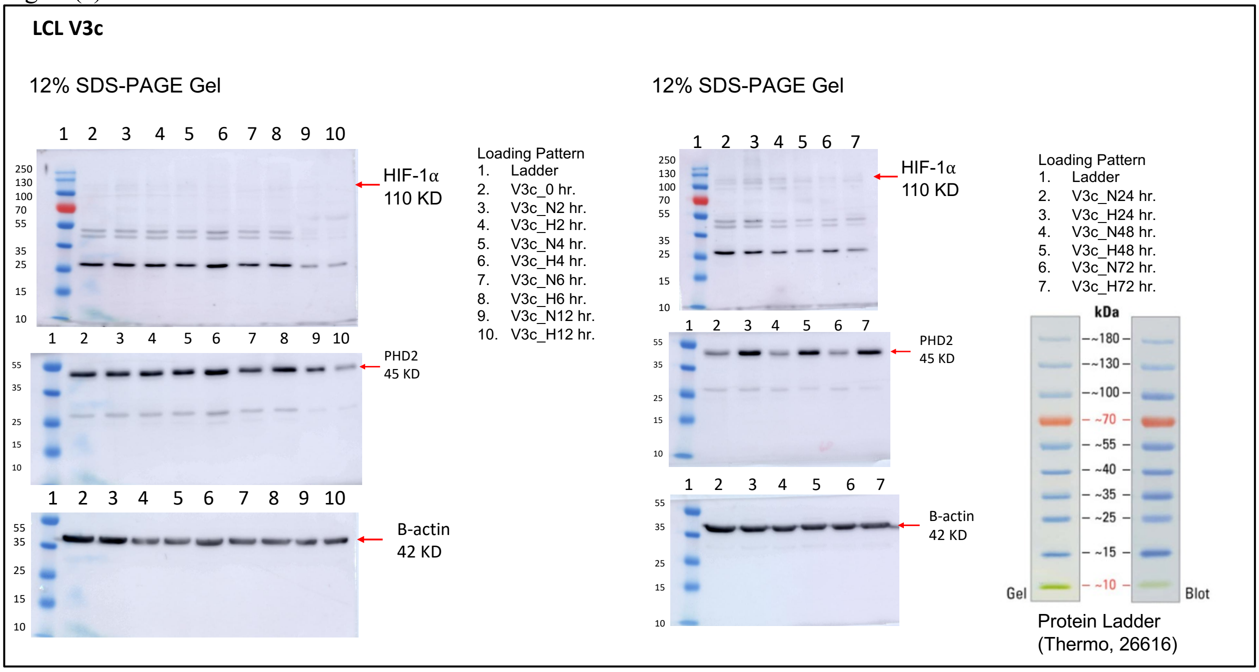

Fig S1(d)

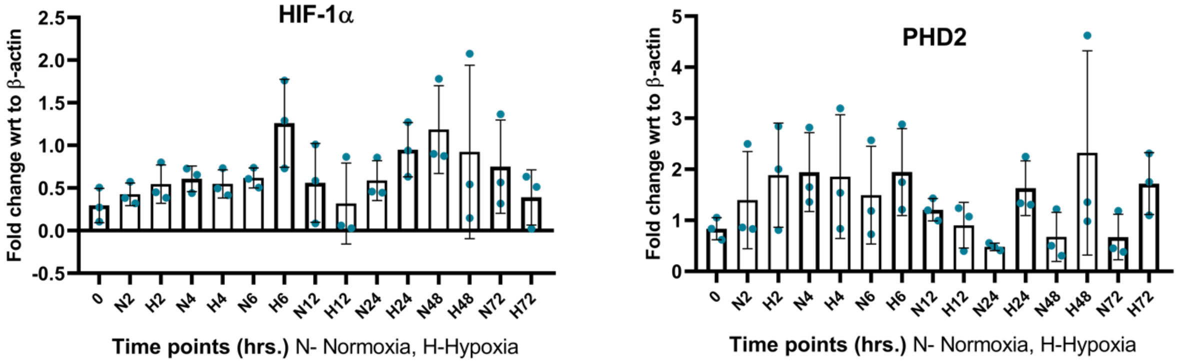


Fig S1: Time point validations using western blotting experiments at selected time points (0, 2, 4, 6, 12, 24, 48, and 72 hrs.). Figure S1(a), Kapha LCL (V1a), S1(b), Pitta LCL (V2c), S1(c), Vata LCL (V3c). S1(d) Combined densitometry data of western blotting experiments for HIF-1α and PHD2 proteins at all selected time points.

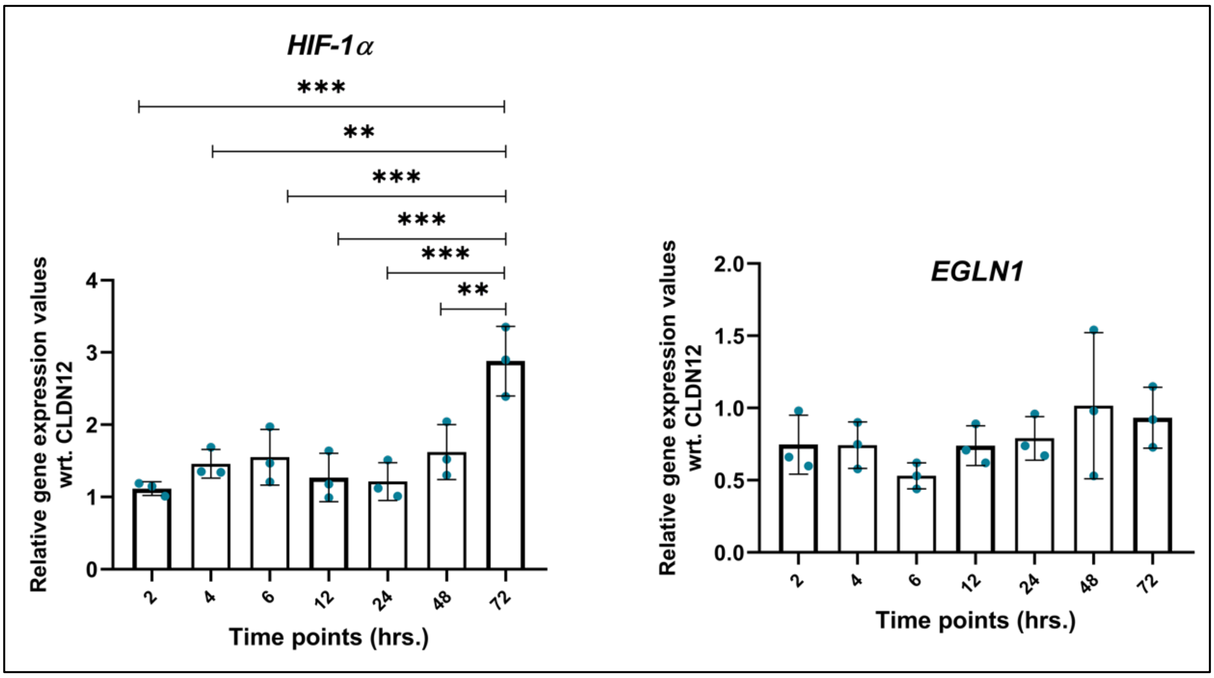


Fig S2: qRT-PCR data for hypoxia generation using 0.2% oxygen and time point validation at selected time points (2, 4, 6, 12, 24, 48, and 72 hrs.). HIF-1α and EGLN1 levels were checked using extreme Prakriti LCLs (one cell line from each Prakriti) treated for hypoxia time points in three biological replicates, N3, n3.

### Baseline comparisons: Ingenuity Pathway Analysis of canonical pathways

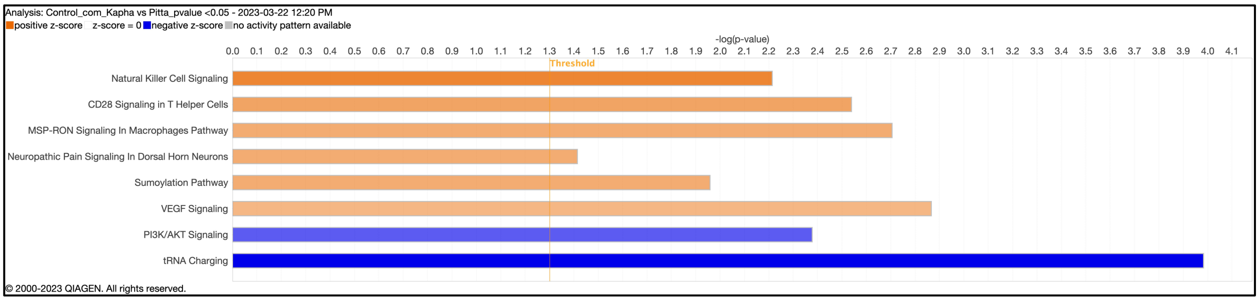


Fig S3: Baseline comparisons (0 hrs.) of Kapha and Pitta using canonical pathways of IPA

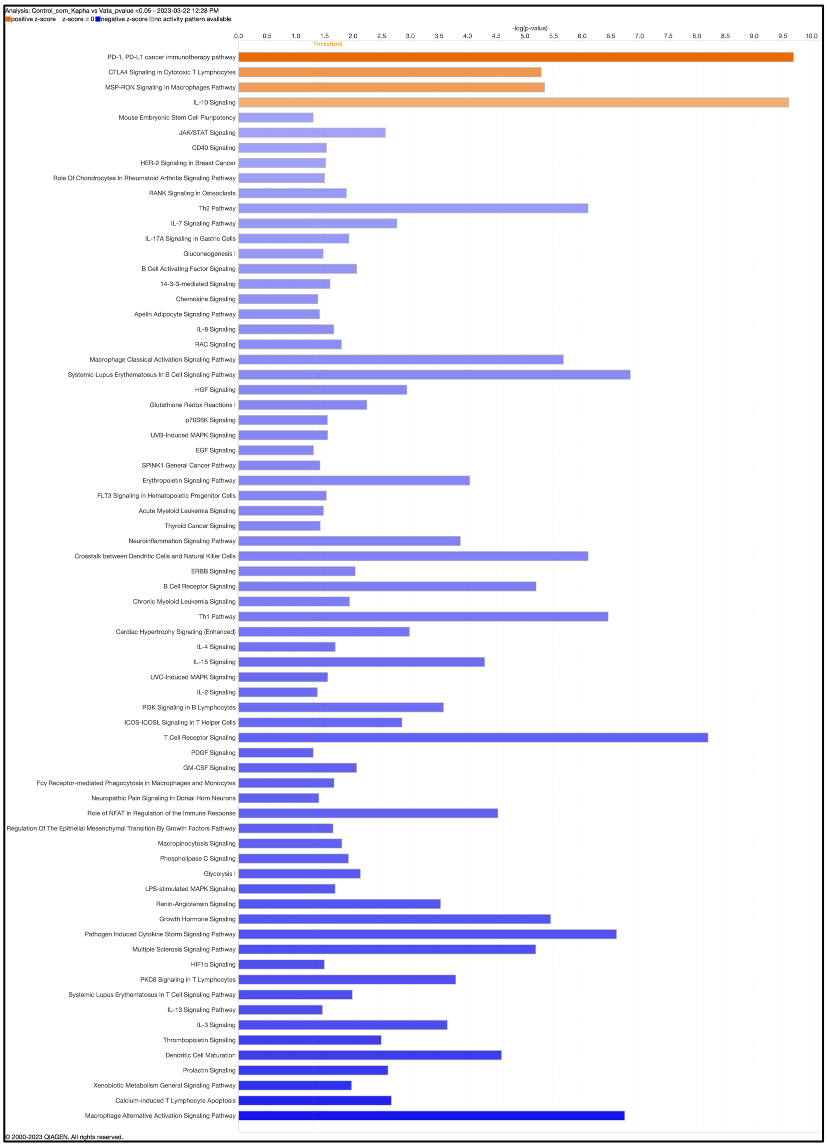


Fig S4: Baseline comparisons (0 hrs.) of Kapha and Vata using canonical pathways of IPA

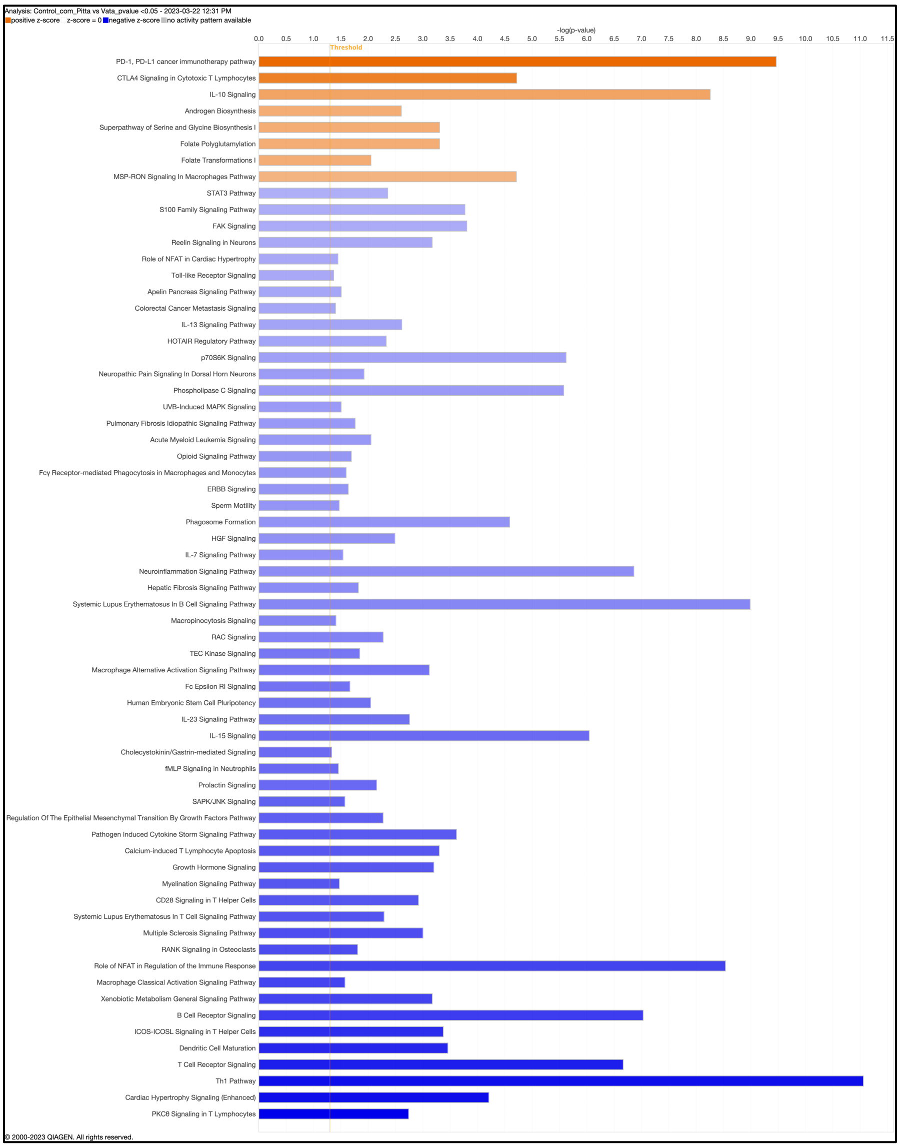


Fig S5: Baseline comparisons (0 hrs.) of Pitta and Vata using canonical pathways of IPA

GSEA-KEGG pathway analysis of pre-ranked gene list

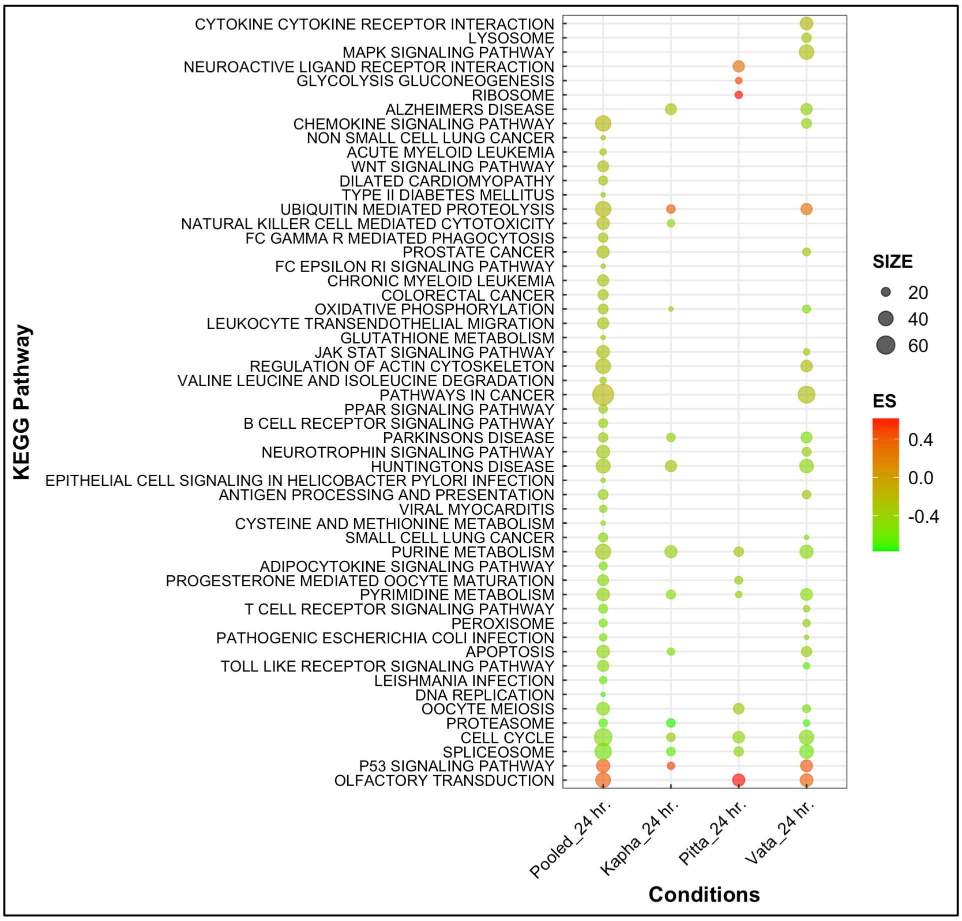


Fig S6: KEGG pathway analysis using GSEA after 24 hrs. of hypoxia.

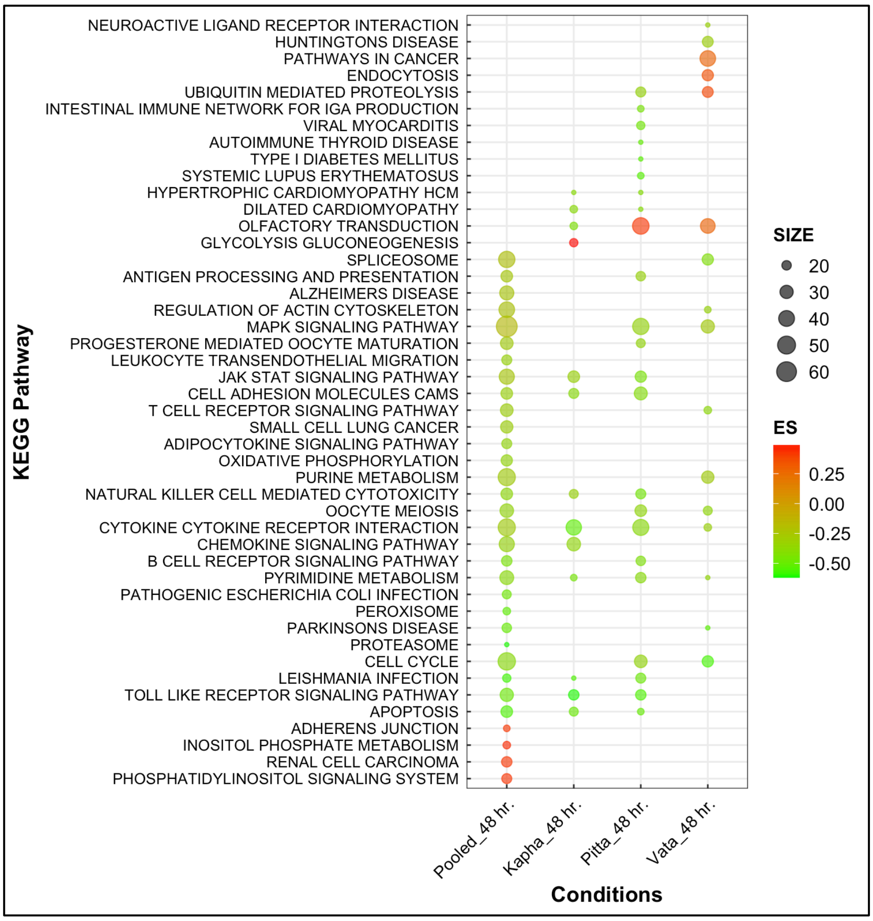


Fig S7: KEGG pathway analysis using GSEA after 48 hrs. of hypoxia.

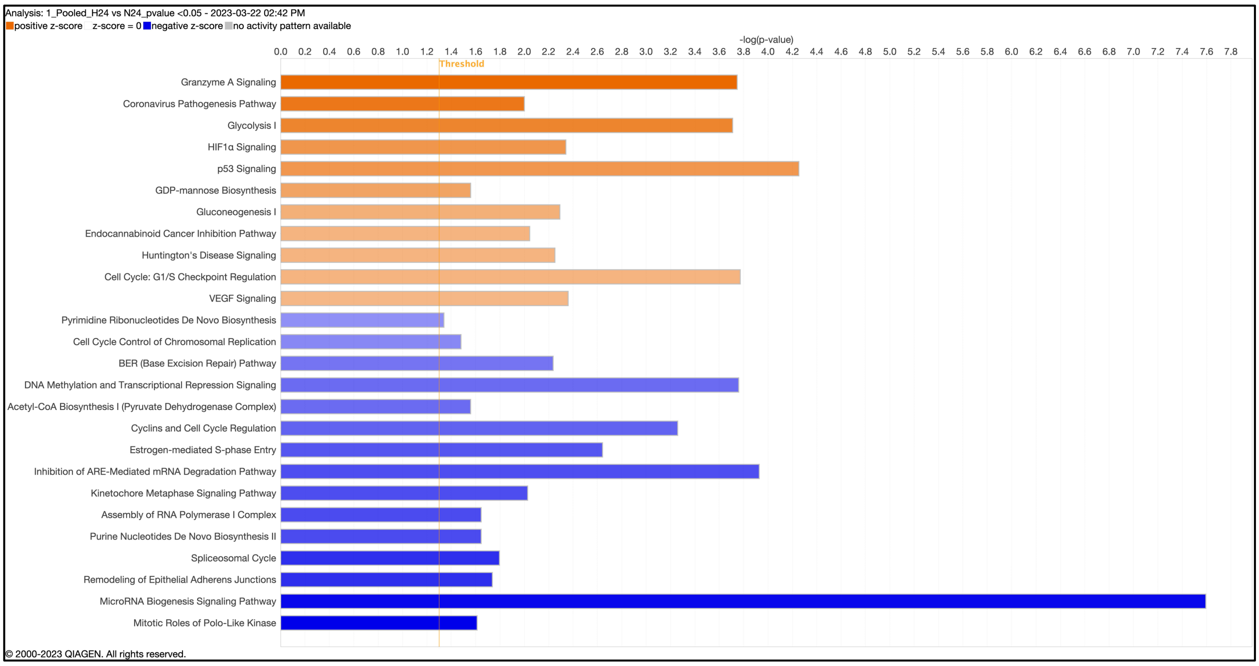


Fig S8: Pooled data enrichment of canonical pathways after 24 hrs. using IPA

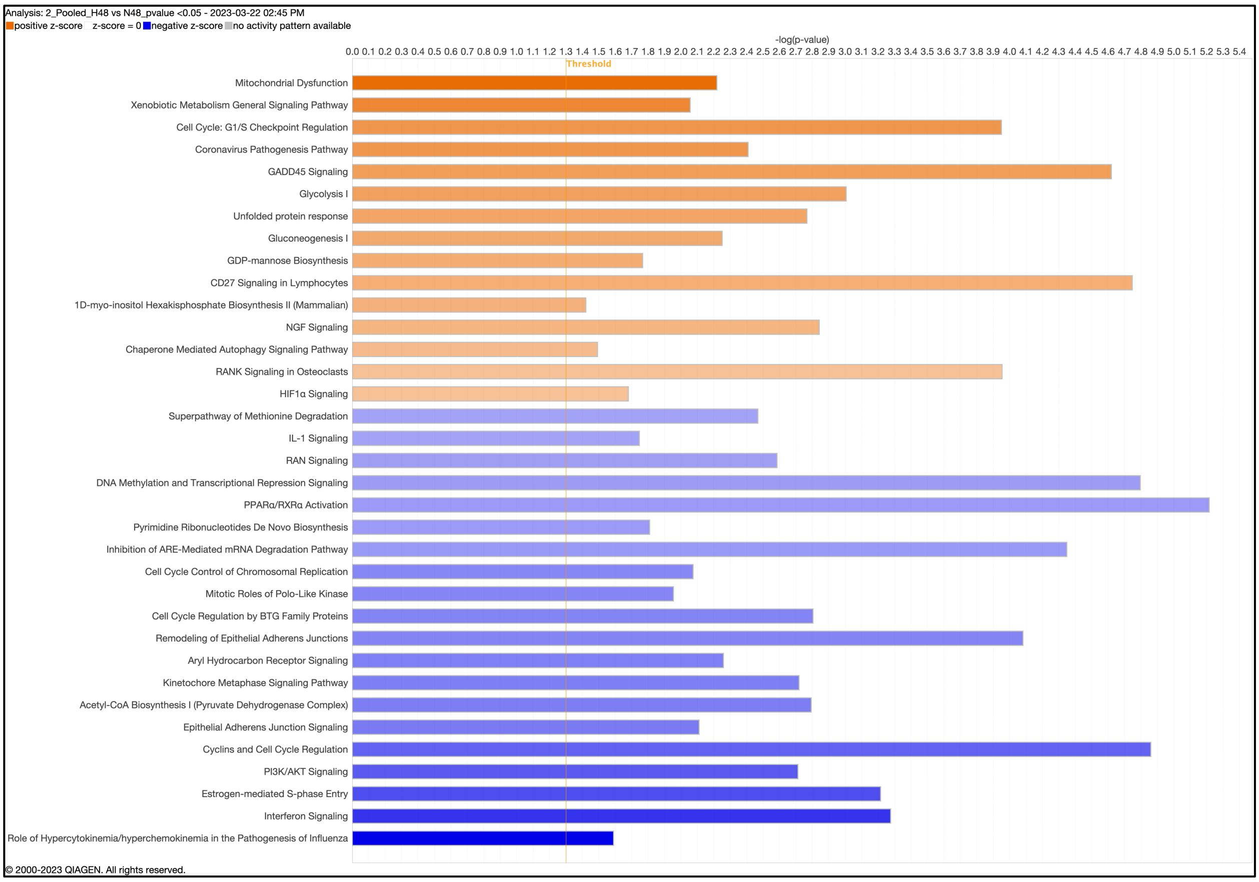


Fig S9: Pooled data enrichment of canonical pathways after 48 hrs. using IPA

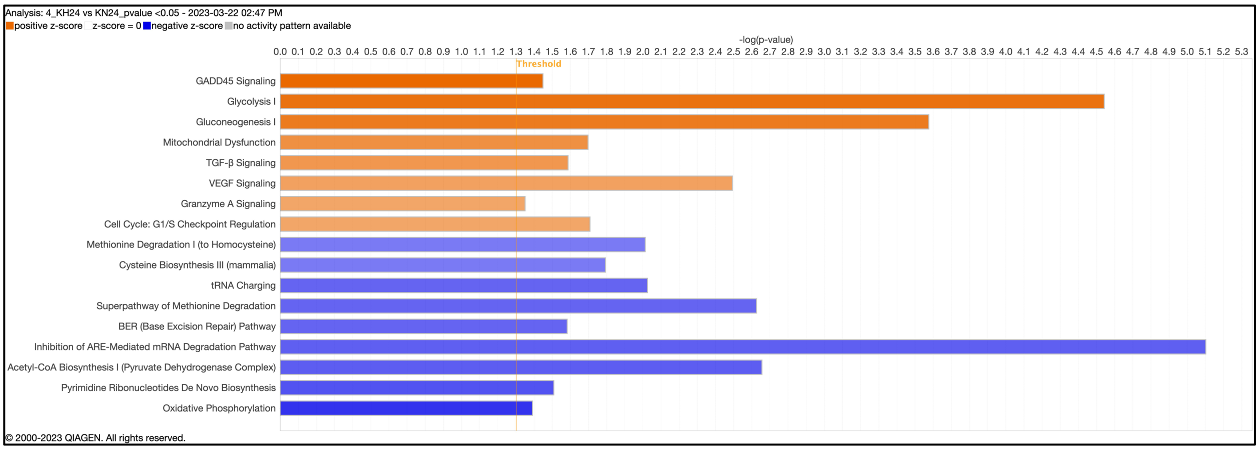


Fig S10: Kapha data enrichment of canonical pathways after 24 hrs. using IPA

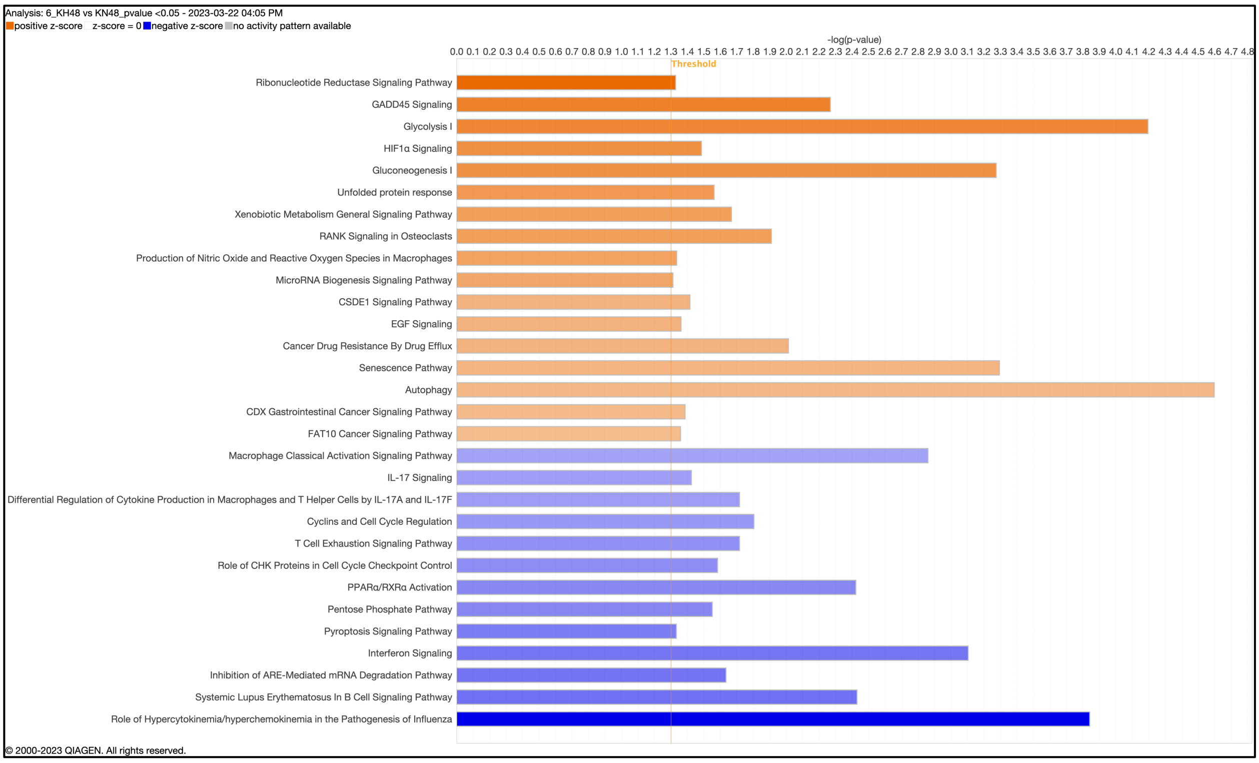


Fig S11: Kapha data enrichment of canonical pathways after 48 hrs. using IPA

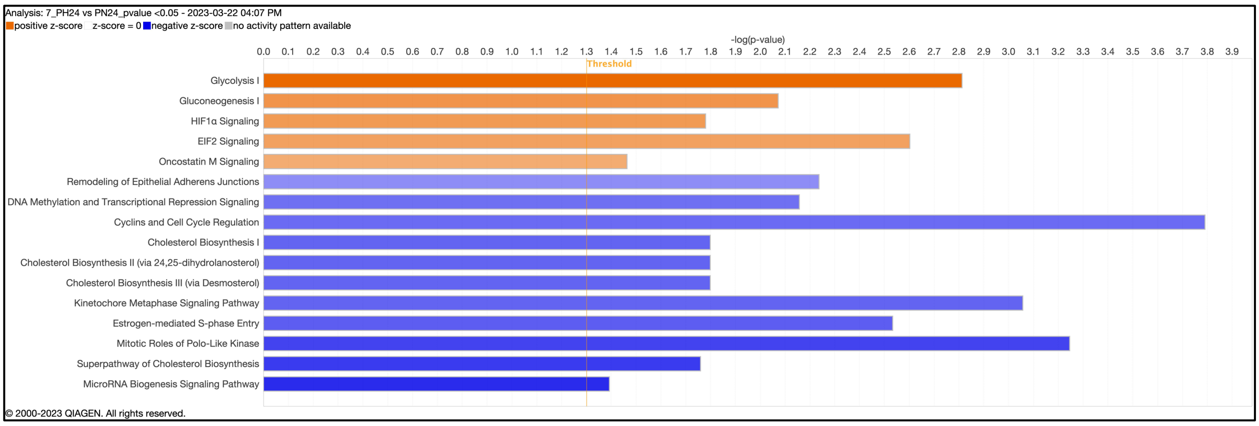


Fig S12: Pitta data enrichment of canonical pathways after 24 hrs. using IPA

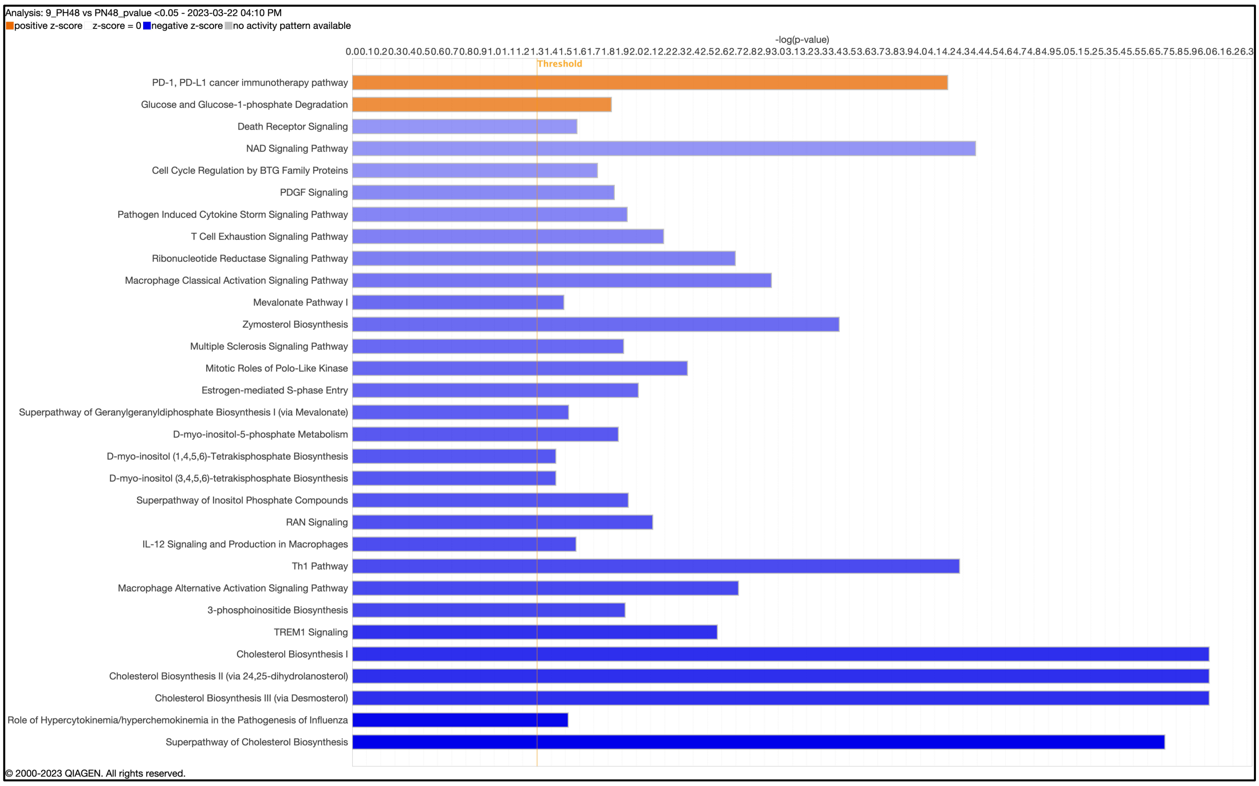


Fig S13: Pitta data enrichment of canonical pathways after 48 hrs. using IPA

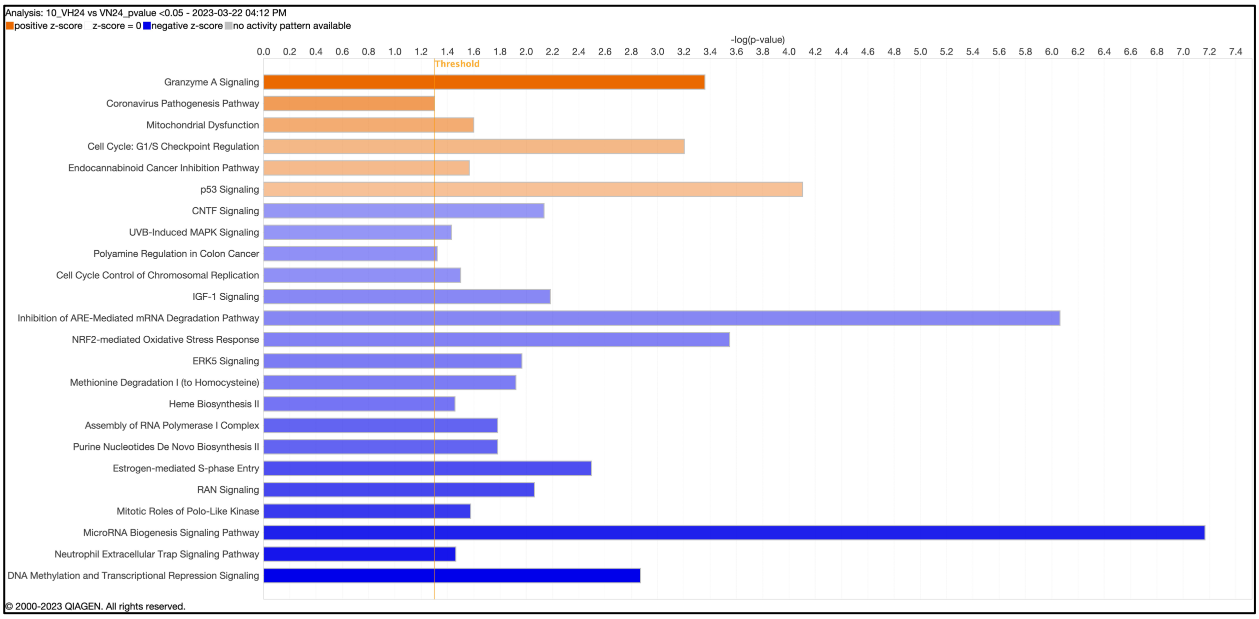


Fig S14: Vata data enrichment of canonical pathways after 24 hrs. using IPA

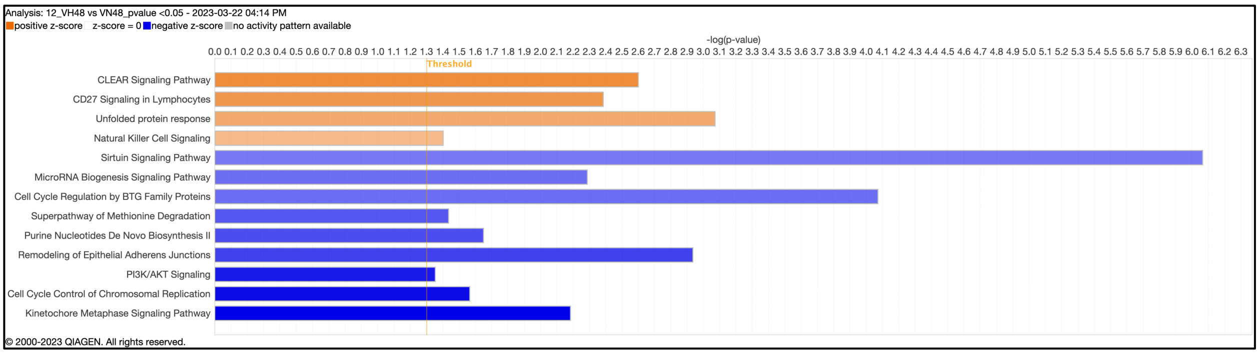


Fig S15: Vata data enrichment of canonical pathways after 48 hrs. using IPA

| Canonical pathways | Pooled | | Kapha | | Pitta | | Vata | |
|---|---|---|---|---|---|---|---|---|
| | 24 | 48 | 24 | 48 | 24 | 48 | 24 | 48 |
| HIF-1a | ↑ | ↑ | | ↑ | ↑ | | | |
| p53 signaling | ↑ | | | | | | ↑ | |
| Glycolysis | ↑ | ↑ | ↑ | ↑ | ↑ | | | |
| Gluconeogenesis | ↑ | ↑ | ↑ | ↑ | ↑ | | | |
| Mitochondrial dysfunction | | ↑ | ↑ | | | | ↑ | |
| Oxidative phosphorylation | ↓ | | ↓ | | | | | |
| Cholesterol biosynthesis | | | | | ↓ | ↓ | | |
| Nucleotide biosynthesis | ↓ | | ↓ | | | | ↓ | ↓ |
| UPR | | ↑ | | ↑ | | | | ↑ |
| GADD45 signaling | | ↑ | ↑ | ↑ | | | | |
| Granzyme A signaling | ↑ | | ↑ | | | | ↑ | |
| Interferon signaling | | | | ↓ | | ↓ | | |
| Cyclin & Cell cycle regulation | ↓ | | | ↓ | ↓ | ↓ | ↓ | |
| Cell cycle checkpoint regulation | ↑ | | ↑ | | | | ↑ | |
| TGF-β | ↑ | | ↑ | | | | | |
| VEGFA | | | ↑ | | | | | |

Fig S16: Summary of IPA canonical pathways in pooled and in all comparisons after 24 and 48 hrs. of hypoxia

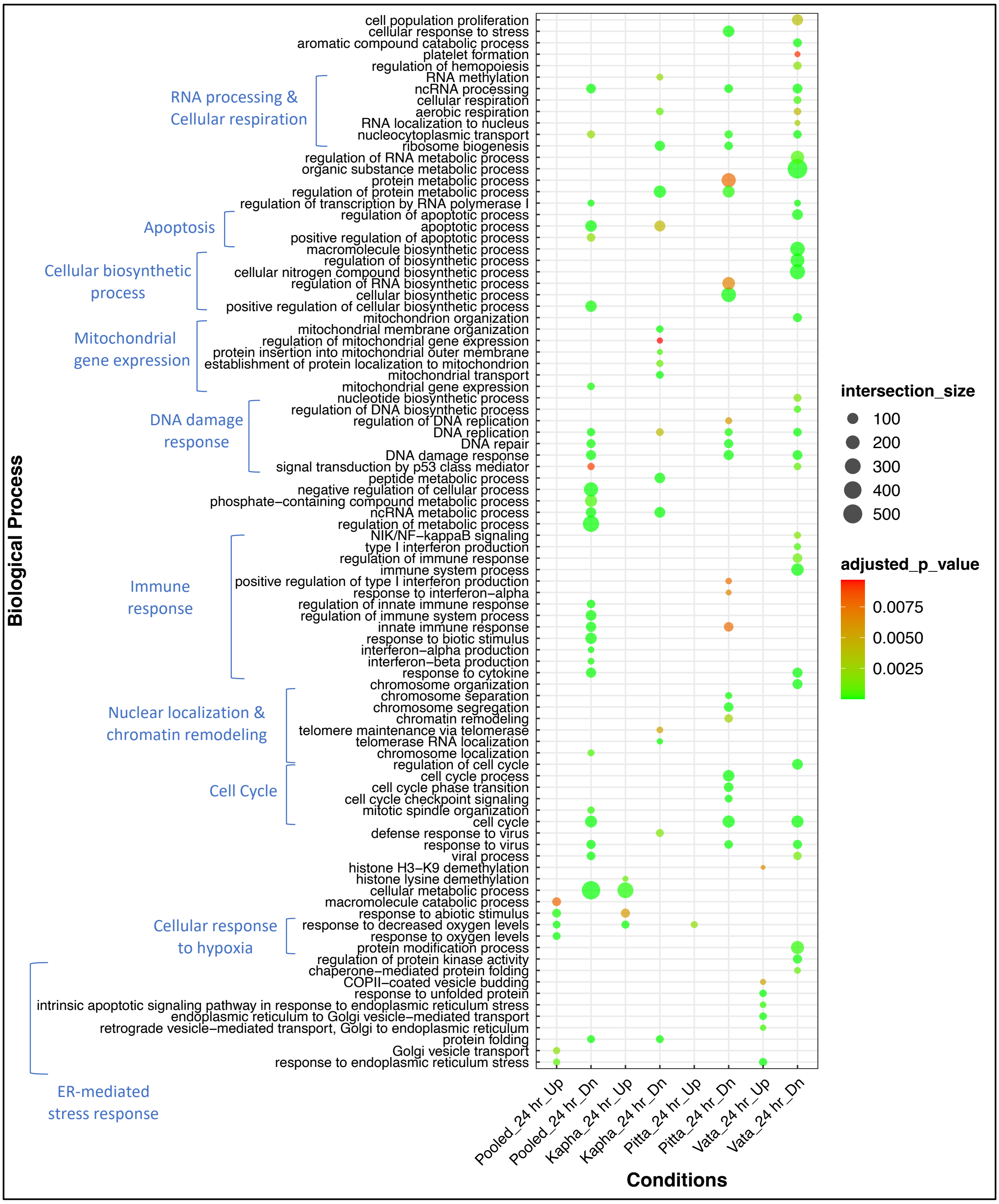


Fig S17: Gene enrichment biological processes using DEG after 24 hrs. with 1.5 fold (>1.5 and <-1.5 FC) using Gene profiler

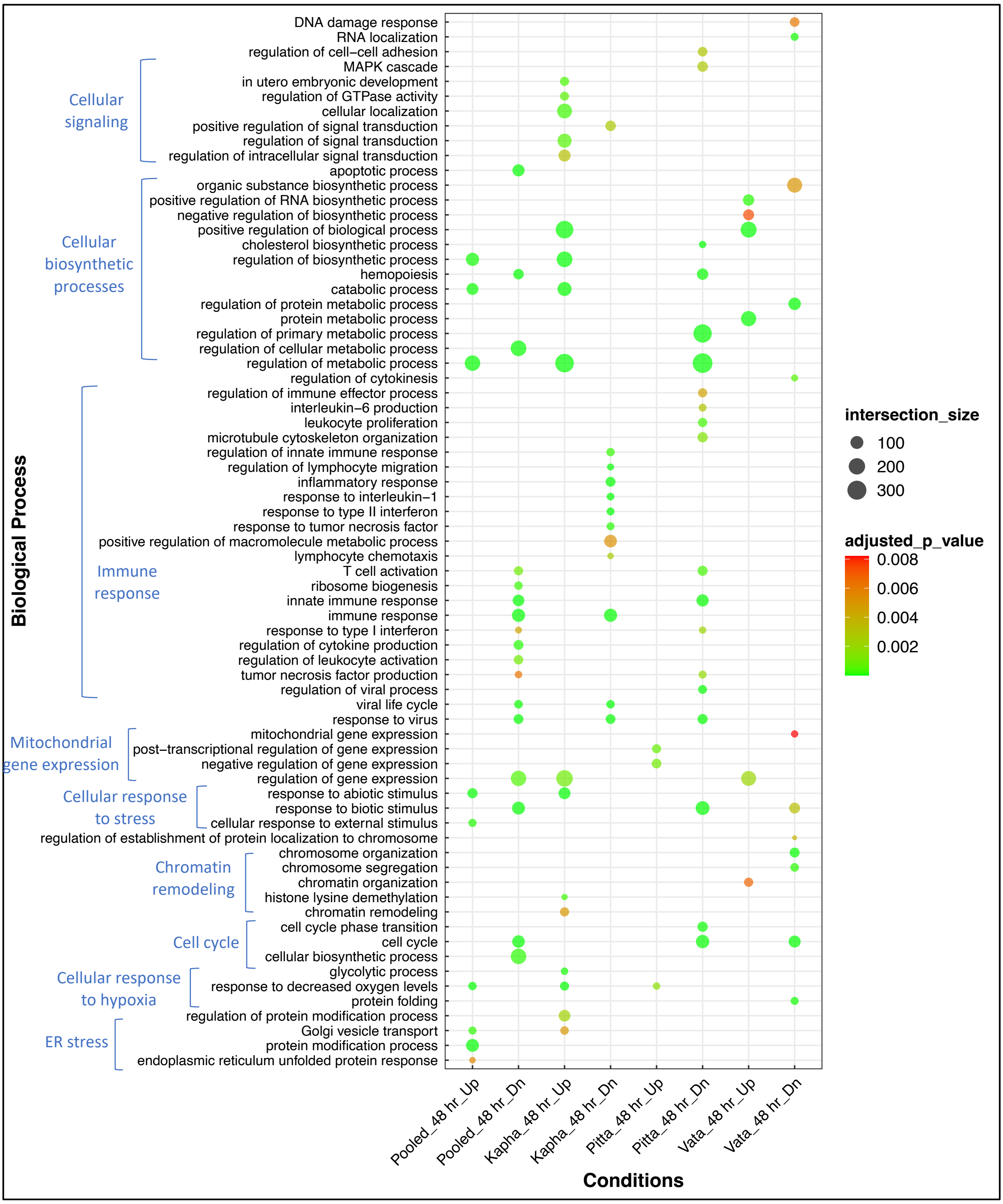


Fig S18: Gene enrichment biological processes using DEG after 48 hrs. with 1.5 fold (>1.5 and <-1.5 FC) using Gene profiler

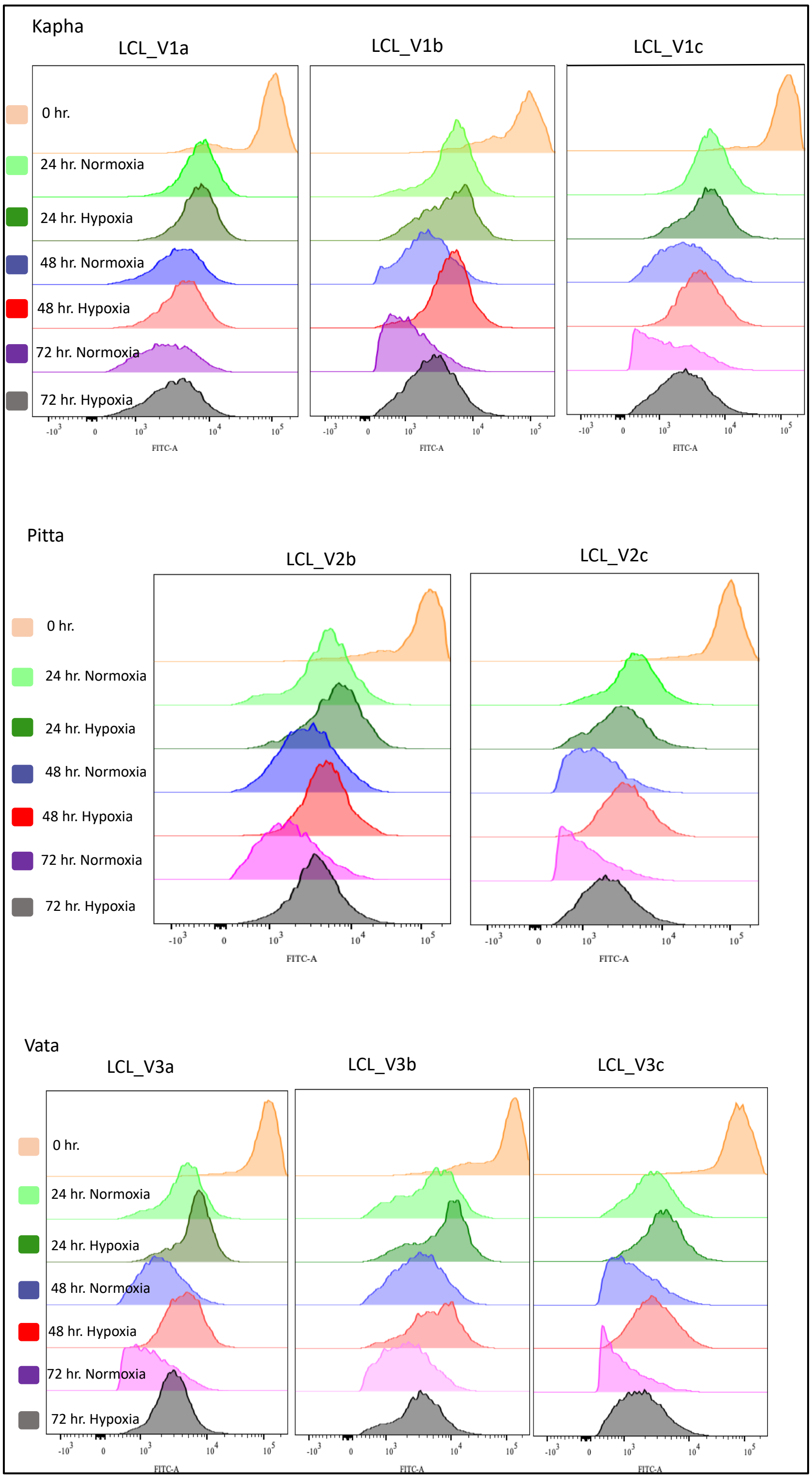


Fig S19: Cell proliferation after hypoxia treatment in Kapha, Pitta, and Vata using CFSE assay for selected time (24, 48, and 72 hrs.) points.
